# Proteomic Mapping of Intercellular Synaptic Environments via Flavin-Dependent Photoredox Catalysis

**DOI:** 10.1101/2022.09.29.510150

**Authors:** Tyler J. Bechtel, Jayde M. Bertoch, Aleksandra K. Olow, Margaret Duich, Cory H. White, Tamara Reyes-Robles, Olugbeminiyi O. Fadeyi, Rob C. Oslund

## Abstract

Receptor-ligand interactions play essential signaling roles within intercellular contact regions. This is particularly important within the context of the immune synapse where protein communication at the surface of physically interacting T cells and antigen-presenting cells regulate downstream immune signaling responses. To identify protein microenvironments within immunological synapses, we combined a flavin-dependent photocatalytic labeling strategy with quantitative mass spectrometry-based proteomics. Using α-PD-L1 or α-PD-1 single-domain antibody (VHH)-based photocatalyst targeting modalities, we profiled protein microenvironments within the intercellular region of an immune synapse-forming co-culture system. In addition to enrichment of both PD-L1 and PD-1 with either targeting modality, we also observed enrichment of both known immune synapse residing receptor-ligand pairs and surface proteins, as well as previously unknown synapse residing proteins.

## Introduction

Many important protein-mediated biological processes occur at the cell surface through cis and trans protein interactions to drive intracellular signal transduction and communication between neighboring cells (**Figure 1a**)^1^. For example, physical cellular association between T cells and antigen-presenting cells (APCs) involve key protein-mediated interactions that include T cell receptors (TCRs) and peptide-MHC complexes (pMHCs), adhesion molecules, and costimulatory/checkpoint proteins^2,3^. How these proteins arrange and engage within these intercellular environments ultimately drive T cell activity with physiological and pathophysiological consequences. For example, transcellular interactions between immune checkpoint ligand PD-L1, overexpressed on tumor cells, and its receptor PD-1, on T cells, results in T cell inactivation that hinders T cell-mediated killing of tumors^4^. Elucidation of this interaction has led to the development of PD-1/PD-L1 immunotherapies, highlighting the importance of understanding immune synapse protein environments. To this end, technologies that unbiasedly assess proteins within these intercellular microenvironments can play an important role in providing a more comprehensive picture of synaptic protein landscapes (**Figure 1b**). Ideally, an approach that leverages the use of therapeutic modalities that effectively target these intercellular regions would open opportunities for profiling synaptic protein microenvironments in a high-resolution manner.

**Figure 1.**
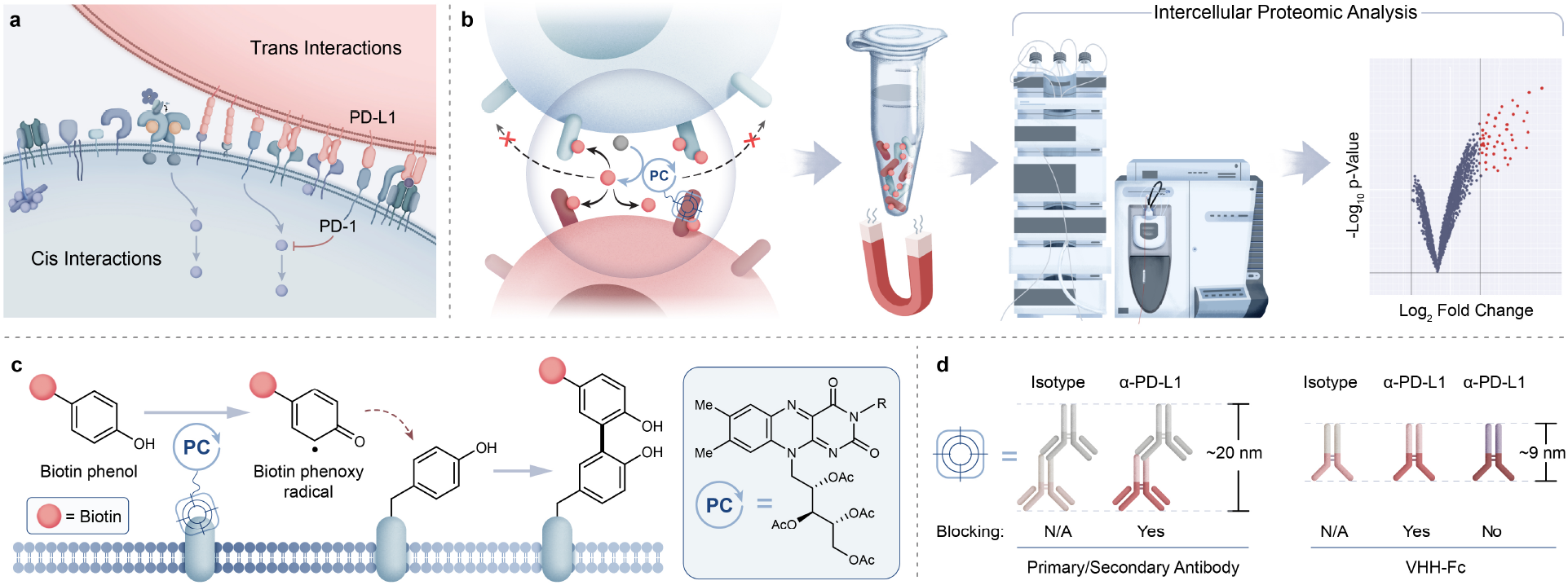
Intercellular photocatalytic protein microenvironment labeling. **a)** Simplified diagram highlighting cis and trans surface protein interactions (such as PD-1 and PD-L1) and their crucial roles in cell signaling. **b)** Schematic depicting targeted photocatalytic proximity labeling at cell-cell interfaces followed by downstream enrichment and identification by MS-based proteomic analysis. **c)** Riboflavin tetraacetate (RFT) is a visible light-activatable photocatalyst capable of generating phenol-based reactive tags that covalently label electron-rich amino acids. **d)** PD-L1 targeting modalities of varying sizes and blocking properties may be used to direct the photocatalyst to PD-L1/PD-1 interacting microenvironments.

Proximity labeling strategies such as BioID^5^, APEX^6^, and TurboID^7^ have been developed that rely on the use of an engineered ligase or peroxidase enzyme fused to a protein of interest to generate reactive biotin-containing species to tag neighboring protein environments. These methods have been extensively used to proteomically profile subcellular compartments^8–10^, as well as cell-cell interfaces^11,12^. While these approaches have been transformative in identifying protein networks within subcellular regions, the enzyme dependency of these technologies may perturb the overall physical properties of the targeting therapeutic modality as well as limit the ability for temporal control of the labeling reaction. Also, the requirement for cell engineering hinders utility within cell types that are not amenable to recombinant expression. Other emerging proximity labeling methods include the use of a two-step photooxidation process wherein enzymes or fluorescent dyes are activated to first generate singlet oxygen species for promiscuous protein oxidation followed by a second step condensation-or disulfide-based trapping of oxidized protein residues^13–16^. Ultimately, despite the many successful efforts to profile a wide range of cellular regions using proteomic-based proximity labeling, implementation of a direct labeling technology within intercellular environments remains a challenge.

Photocatalytic-based methods offer an attractive alternative to the above approaches through their relatively much smaller size compared to enzymes, their exquisite spatiotemporal control, and their ability to induce direct activation of a biotin probe for targeted surface protein labeling^17–19^. Indeed, such efforts have been deployed for covalent labeling and mass spectrometry (MS)-based identification of cis-interacting protein environments on mammalian cell surfaces^20–23^. Photocatalytic-based approaches are also expanding into more complex cellular environments as demonstrated through our recently disclosed photocatalytic cell tagging (PhoTag) technology^24^. This method exploits visible light-activation of flavin co-factors to generate phenoxy radicals^21^ (**Figure 1c**) for selective tagging of physically associated cellular interfaces to facilitate downstream transcriptome analysis of cell-cell interactions. Given the importance of identifying proteins that reside within synaptic microenvironments, we envisioned that this flavin photocatalytic system would be well suited for extending beyond current photocatalyst-based profiling activities in monoculture systems and enable the proteomic analysis of intercellular protein interactions. Here we describe the development of a robust MS-based intercellular proteomics platform that leverages the use of VHH-based photocatalyst conjugates to capture and identify synaptic protein microenvironments (**Figure 1b**).

## Results and Discussion

To identify an optimal targeting modality for profiling the immune synapse, we began our studies by first investigating the use of a primary/secondary antibody or a blocking/non-blocking VHH-Fc photocatalyst conjugate system to label PD-L1 microenvironments on the surface of B cells in monoculture (**Figure 1c**)^24^. Accordingly, we site-selectively functionalized a blocking α-PD-L1 VHH-Fc that disrupts the PD-1/PD-L1 interaction with riboflavin tetraacetate (RFT) (**Supplementary Figure 1a**) for PD-L1-targeted proximity labeling. JY B cells overexpressing PD-L1 (hereafter referred to as JY PD-L1 cells) were incubated with α-PD-L1 VHH-Fc-RFT followed by the addition of biotin phenol (**Supplementary Figure 1b**) and irradiated with visible blue light (450 nm). Western blot analysis showed light- and time-dependent biotinylation compared to an isotype control antibody and confocal imaging analysis confirmed that these labeling events were selective for the cell surface (**Figure 2a** and **Supplementary Figure 2**). Next, biotinylated proteins were enriched on streptavidin magnetic beads and identified by tandem mass tag (TMT)-based quantitative LC-MS/MS proteomic analysis (**Figure 2b** and **Supplementary Table 1**). Targeted labeling with α-PD-L1 VHH-Fc-RFT resulted in PD-L1 being the most highly enriched protein with significant enrichment of other immune relevant surface proteins such as HLA class I and II molecules, CD40, and ICAM protein family members, highlighting the ability of this technology to identify proteins on the surface of B cells in a targeted and proximity-based fashion (**Figure 2b**). In an analogous experiment, a primary/secondary antibody system, comprised of a commercially available blocking α-PD-L1 primary antibody and a goat anti-mouse secondary antibody non-site selectively labeled with RFT, yielded similar results by western blot and confocal microscopy (**Supplementary Figure 3**). Likewise, proteomic analysis of the two-antibody system showed strong enrichment for PD-L1 as well as several other proteins similarly identified by VHH-Fc (**Supplementary Figures 4a and 5** and **Supplementary Table 2**).

**Figure 2.**
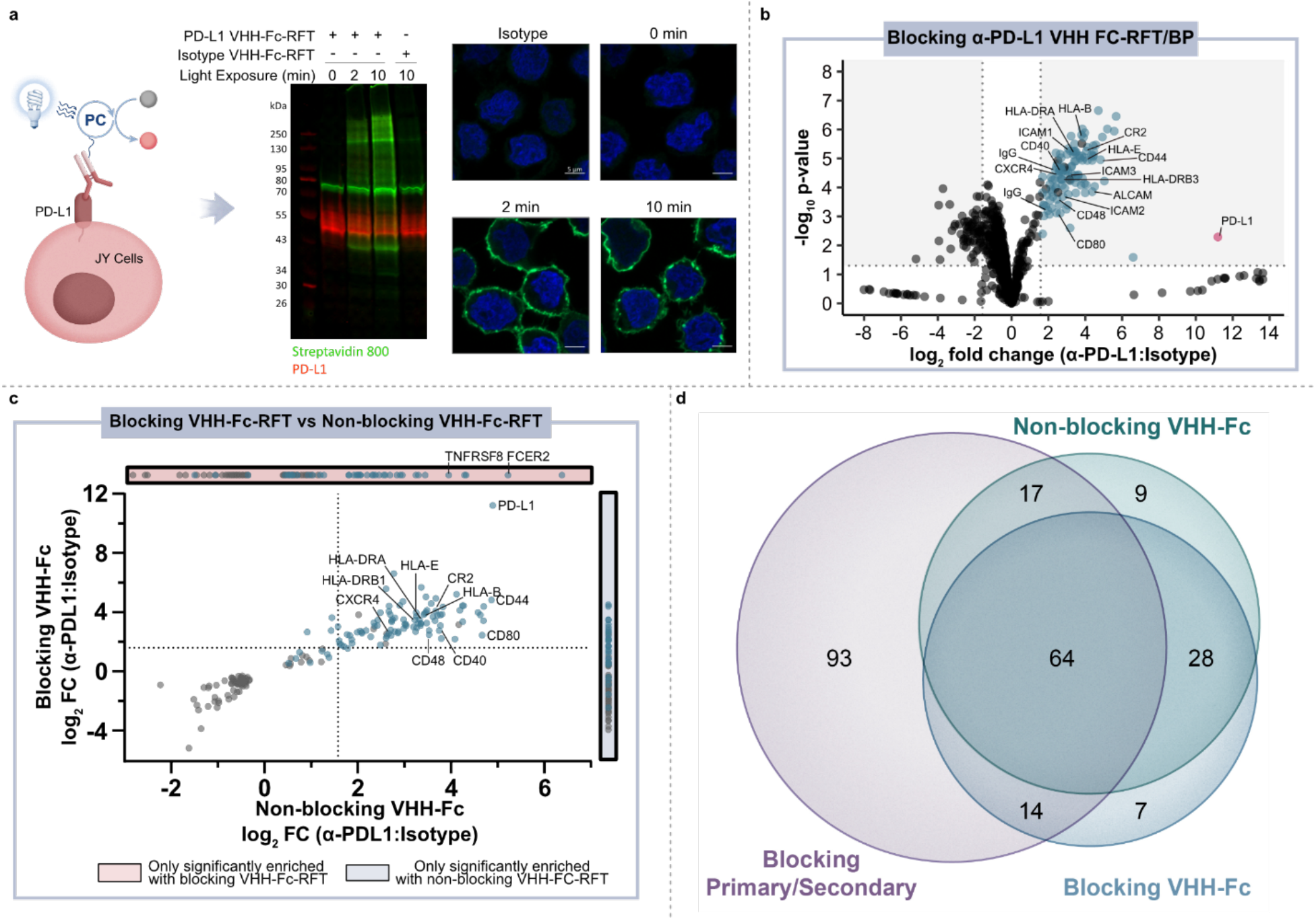
Photocatalytic proximity labeling of PD-L1 microenvironments on the surface of B cells. **a)** Schematic depicting α-PD-L1 VHH-Fc-RFT targeted labeling on the surface of JY PD-L1 cells and western blot and immunofluorescent microscopy analysis of labeling events. Biotinylation levels (green) increase as a function of blue-light irradiation time in PD-L1 targeted samples, but not isotype controls. Nuclei are labeled with Hoechst stain (blue) and scale bars indicate 5 μm. **b)** Volcano plot of statistical significance vs. fold-enrichment for α-PD-L1 VHH-Fc targeted vs isotype-targeted biotinylation on JY PD-L1 cells following 2 minutes of blue-light irradiation. Significantly enriched cell-surface proteins (p-value < 0.05 and log_2_FC > 1.58) are indicated as blue dots and PD-L1 is indicated as a red dot (n = 3 experiments). **c)** Quadrant plot comparing enrichment of proteins in blocking VHH-Fc and non-blocking VHH-Fc PD-L1 targeted datasets. Only proteins that are significantly enriched (p-value < 0.05) in both datasets are displayed. Cell-surface-localizing proteins, determined by Uniprot^35^ and the Surfaceome^36^, are indicated with a blue dot. **d)** Venn diagram depicting the overlap of significantly enriched proteins (p value <0.05 and log_2_FC > 1.58) identified with α-PD-L1 targeting modalities that include a blocking primary/secondary antibody-RFT, blocking VHH-Fc-RFT, and non-blocking VHH-Fc-RFT.

Encouraged by the targeted labeling results of our PD-L1 blocking modalities, we next explored the use of a non-blocking α-PD-L1 VHH-Fc-RFT that does not disrupt PD-1/PD-L1 interactions and serves as a spectator within the PD-L1 microenvironment. As expected, PD-L1 was highly enriched along with other surface proteins that included a well-known PD-L1 *cis* interactor, CD80^25^ (**Supplementary Figure 4b** and **Supplementary Table 3**). Enrichment of this *cis* interactor resulted from the use of the non-blocking VHH that, in addition to lacking PD-1/PD-L1 blocking capacity, does not block PD-L1/CD80 interactions^24^. Differences in protein enrichment between blocking and non-blocking α-PD-L1 VHH-Fc targeting modalities were analyzed using a quadrant plot that compares the fold change enrichment for statistically significantly enriched proteins identified in blocking and/or non-blocking proximity labeling experiments (**Figure 2c**). While many proteins, such as PD-L1, CD44, ALCAM, CR2, CXCR4, ICAM1, and HLA molecules were highly enriched by both VHH modalities, CD80 enrichment was much more pronounced with the non-blocking VHH-Fc compared to the blocking VHH-Fc which blocks both PD-1/PD-L1 and PD-L1/CD80 interactions.

Venn diagram analysis of the enriched protein hits across all three PD-L1 targeting modalities (primary/secondary antibody, blocking VHH, non-blocking VHH) on JY PD-L1 monoculture cells revealed shared overlap of 64 proteins (**Figure 2d and Supplementary Table 4**), many of which have been implicated in PD-L1 biology. For instance, ADAM10 and ADAM17 have been reported to directly cleave the PD-L1 surface domain leading to increased checkpoint inhibitor resistance^26^. EPHA3 has been recently shown to physically interact with PD-L1 in a checkpoint inhibitor (atezolizumab) dependent fashion^27^. Other proteins such as CR2^28^ and HLA molecules^29,30^ have been observed to co-express with PD-L1. Interestingly, inhibition of CXCR4^31^ and SEMA4D^32^ as well as a CD40^33,34^ agonist (all proteins enriched from PD-L1 targeting) are being investigated for evidence of preclinical enhancement in combination with PD-1/PD-L1 checkpoint blockade in patients. To the best of our knowledge, the combined identification of CXCR4, SEMA4D and CD40 has not been previously shown to be proximal to PD-L1 outside of our photocatalyst labeling technologies (**Supplementary Figure 6**), highlighting the use of therapeutic-based modalities as an attractive approach for profiling relevant cell surface environments for discovery of novel drug targets. Finally, in further comparison of the RFT-based VHH and primary/secondary antibody systems, we observed a higher overlap of shared proteins between the VHH targeting modalities as well as a much smaller number of total enriched proteins using the VHHs compared to the primary/secondary antibody system (**Figure 2d**). Collectively, these results showcase the high-resolution nature of this technology where targeting modality size (primary/secondary antibody vs. VHH) and/or binding mode (blocking vs. non-blocking) ultimately impact the microenvironment that is captured on the cell surface.

Having showcased this protein labeling technology in a monoculture system, we next explored the feasibility of applying our labeling technology within immune synapse environments. For this experiment, we used a co-culture system comprised of engineered Jurkat T cells (CD8^+^, PD-1^+^) that also express a TCR with specificity for HLA-A02^37^ (hereafter referred to as Jurkat JS86-PD-1 cells) and JY PD-L1 cells (HLA-A02 positive). Co-incubation of these two cell types results in T cell activation as measured by IL-2 production (**Supplementary Figure 7**). For immunosynaptic PD-L1-targeted proximity labeling, JY PD-L1 cells were pre-labeled with the non-blocking α-PD-L1 VHH-Fc-RFT and co-incubated with Jurkat JS86-PD-1 cells for 30 minutes to allow for cell interaction, after which biotin phenol was added and cells were irradiated with blue light. Under these conditions, flow cytometry analysis confirmed *cis* and *trans* labeling of both cell types and confocal microscopy revealed highly selective labeling events within APC-T cell contact sites (**Supplementary Figure 8 and 9**). To confirm that we are not only tagging PD-L1, but also achieving transcellular tagging of the cognate receptor, PD-1, labeled cells were lysed and biotinylated proteins were enriched using streptavidin magnetic beads followed by elution and downstream LC-MS based proteomic analysis (**Figure 3a**). As anticipated, we observed strong enrichment of both PD-L1 and PD-1, (**Figure 3b, middle panel** and **Supplementary Table 5**) confirming the ability of our method to achieve protein labeling on both sides of the immune synapse.

**Figure 3.**
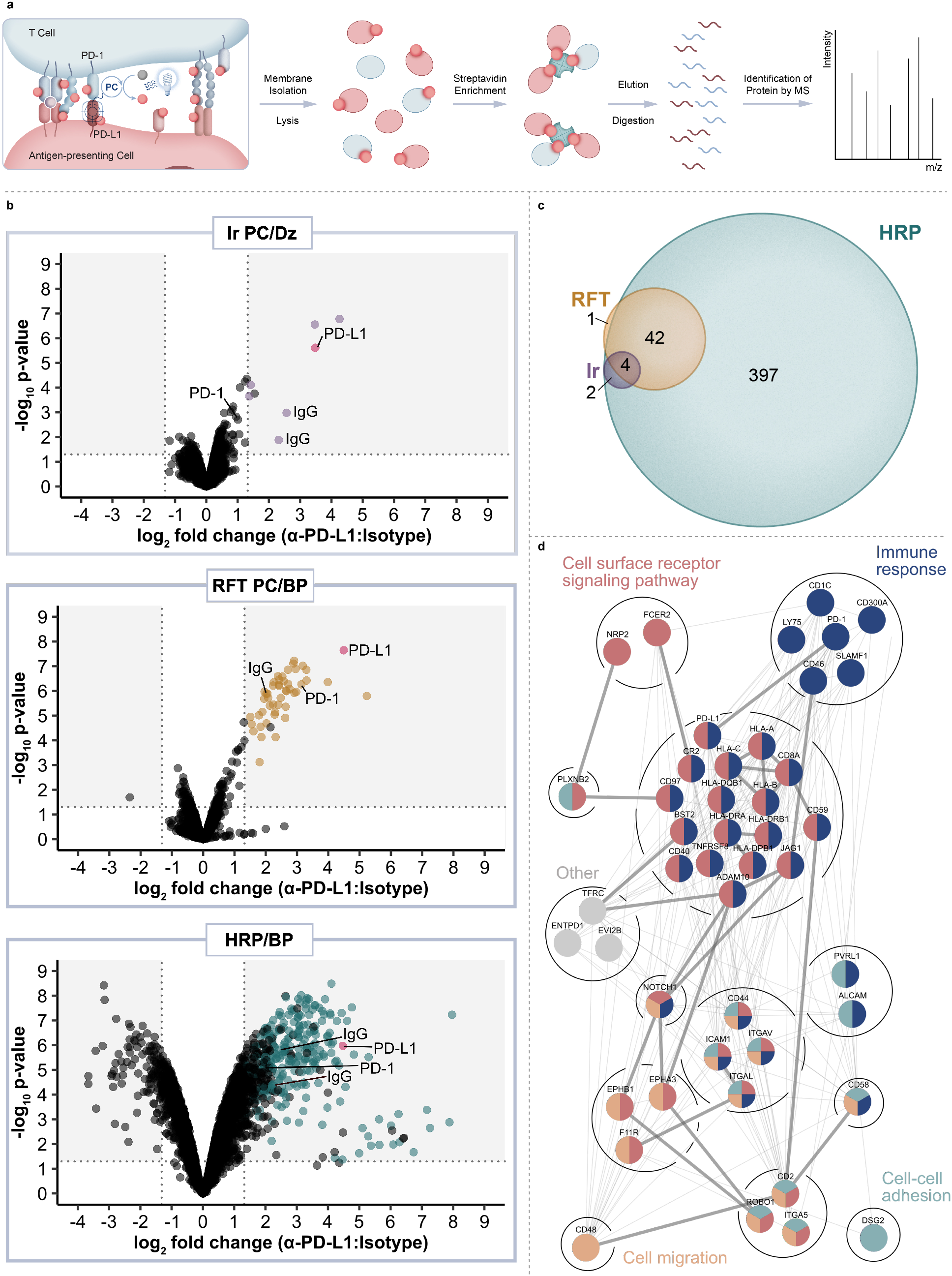
Intercellular proteomic platform for identification of PD-L1 microenvironments within the immunological synapse of Jurkat JS86-PD-1 T cells and JY PD-L1 B cells. **a)** Schematic depicting intercellular photocatalytic proximity labeling within the immunological synapse and downstream proteomic workflow. **b)** Comparison of PD-L1 targeted catalytic labeling technologies including an iridium photocatalyst (purple), RFT (yellow), and horseradish peroxidase (blue), within the immune synapse. Data are represented as volcano plots of statistical significance vs. fold-enrichment for non-blocking VHH-Fc PD-L1 targeted vs isotype-targeted biotinylation. Significantly enriched cell-surface proteins (p-value < 0.05 and log_2_FC > 1.32) are indicated as purple, yellow, or blue dots and PD-L1 is indicated as a red dot (n = 3 experiments). **c)** Venn diagram depicting overlap between Ir PC, RFT, and HRP intercellular proximity labeling approaches. Significantly enriched proteins from each method were included (p-value <0.05 and log_2_FC > 1.32) **d)** STRING^38^ protein interaction network of significantly enriched PD-L1 proximal proteins within the immune synapse (p-value <0.05 and log_2_FC > 1.32). Proteins are further grouped by biological process GO^39^ terms. Coloration indicates membership within broad gene ontology terms and multiple colors indicate nodes within multiple terms. Thick edges represent interaction evidence from experimental evidence while thin edges indicate evidence from other sources.

To compare the ability of other methods to target and enrich PD-L1 and PD-1 within the synapse, we performed a similar labeling experiment using either a peroxidase enzyme that also generates a phenoxy radical for protein labeling or an iridium photocatalyst (Ir PC)-based micromap method that we recently disclosed^20^ for carbene-mediated proximal protein labeling. Accordingly, we conjugated non-blocking α-PD-L1 VHH-Fc to HRP or the iridium photocatalyst for labeling within the JY-Jurkat co-culture system. While proximity labeling via HRP in the presence of biotin phenol resulted in enrichment of PD-L1 and other proteins in similar fashion to RFT-mediated labeling, the overall number of significantly enriched proteins was nearly 10-fold higher with HRP (**Figure 3b, bottom panel, Figure 3c** and **Supplementary Table 6**). Furthermore, PD-1 protein enrichment was much less pronounced with HRP-mediated labeling compared to the RFT system (**Figure 3b, middle and bottom panel**). These results are consistent with previous observations of promiscuous labeling that occurs inside and outside of the synaptic region using HRP^24^ and as further confirmed by confocal imaging within this JY-Jurkat co-culture system (**Supplementary Figure 9**). On the other end of the spectrum, targeted labeling with iridium (**Supplementary Figure 1c**) in the presence of diazirine biotin (**Supplementary Figure 1d**) led to significant enrichment of PD-L1 in similar fashion to RFT but did not strongly enrich other proteins including PD-1 from the T cell surface (**Figure 3b, top and middle panel, Figure 3c** and **Supplementary Table 7**). This observation is supported by the shorter labeling radius of the carbene intermediate that displays a much shorter half-life^20^ than the phenoxy radical from RFT or HRP labeling^17,18^. These combined results demonstrate the enhanced ability of RFT-mediated synaptic labeling to co-enrich PD-L1 and its cognate receptor, PD-1, from synaptic environments compared to HRP and iridium-based micromap methods.

In addition to PD-L1 and PD-1 enrichment in the synapse with the RFT method, we detected other immune synapse ligand-receptor partners that include ICAM-1/LFA-1α, CD2/CD2 ligands (CD48, CD58 and CD59), JAG1/NOTCH1, CD8A/HLA class I molecules as well as other proteins with known roles within the immune synapse (CD40, CD46, CR2, and HLA class II molecules) (**Supplementary Figure 10, Supplementary Tables 5 and 8)**. We also identified several proteins with no known association within the immune synapse including adhesion molecule DSG2, receptor tyrosine kinase EPHA3, semaphorin cell surface receptor PLXNB2, and integrin ITGAV suggesting a potential role for these proteins within T cell-APC interactions **(Supplementary Tables 5 and 8)**. Analysis of all significantly enriched synaptic proteins using the search tool for retrieval of interacting genes (STRING)^38^ revealed mostly predicted (195) versus experimentally annotated (30) protein-protein interactions (**Figure 3d**), further showcasing the utility of our approach to expand on what is known about these protein interactions, particularly within the context of immune synapse environments. Finally, reverse proximity labeling through use of a non-blocking α-PD-1 VHH Fc-RFT within the JY-Jurkat cell system strongly enriched for both PD-1 and PD-L1 (**Supplementary Figure 10 and 11, Supplementary Table 9**). Of the total enriched protein hits from α-PD-1 targeted labeling, 13 proteins overlapped with the PD-L1 targeted list **(Figure 4a)**. As expected, the combined enrichment of proteins from α-PD-1 and/or α-PD-L1 showed GO^39^ term enrichment for immune-related biological processes that includes regulation of the immune response, positive regulation of cytokine production, and heterotypic cell-cell adhesion (**Figure 4b** and **Supplementary Table 10**). In addition, several of the enriched proteins (CD1c, CD40 and LY75) are currently undergoing clinical assessment in combination with PD-1/PD-L1 blockade, while others (CD2, ENTPD1, CD46 and EPHA3) are being evaluated as emerging cancer-based immunotherapies (**Supplementary Table 8)**. Collectively, these observations underscore the robust nature of flavin-based photocatalytic system for capturing functionally relevant protein microenvironments from targeted labeling of either side of the synapse.

**Figure 4.**
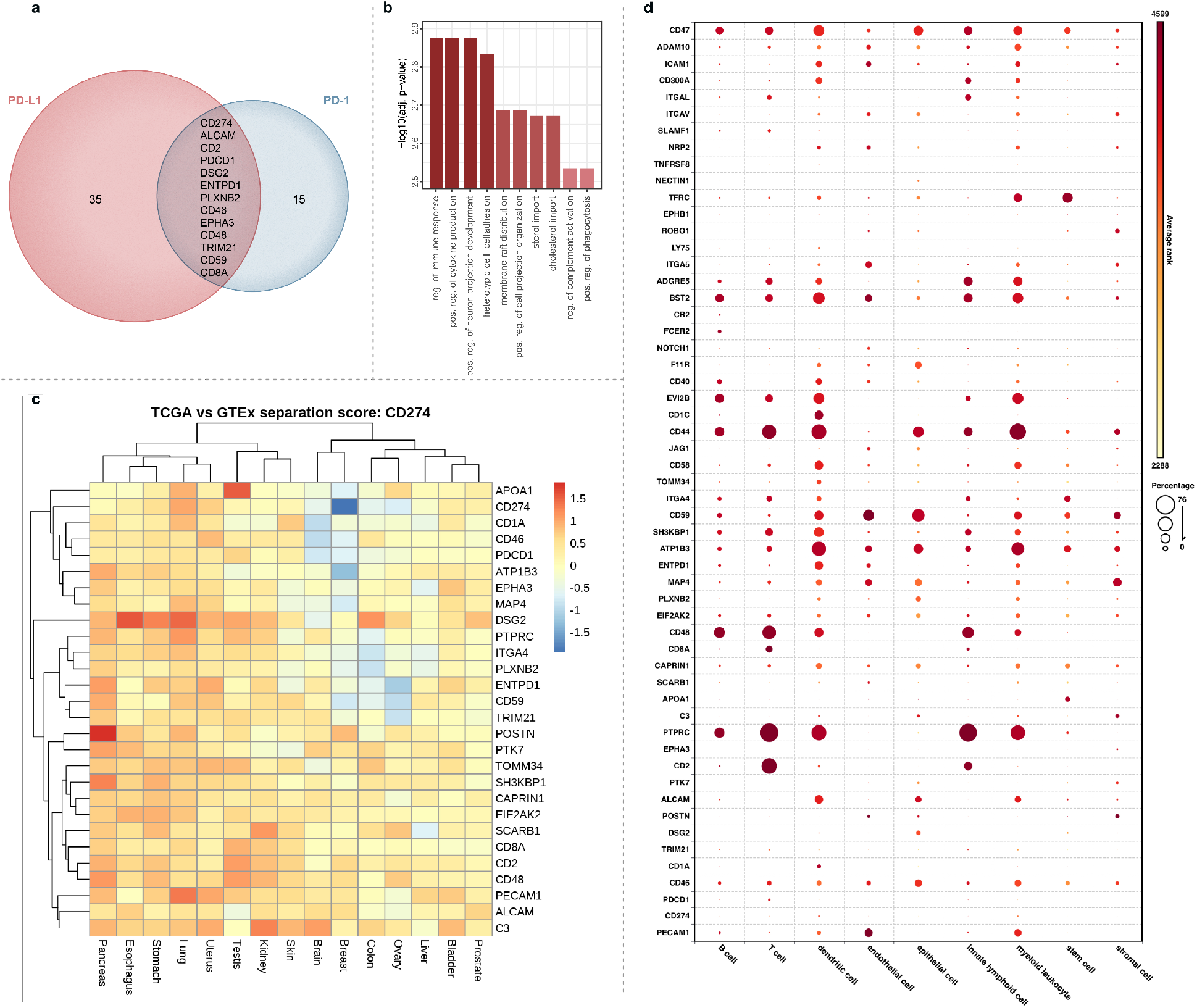
Bioinformatic analysis of significantly enriched proteins from PD-L1 and PD-1 targeted cell-cell interaction environments. **a)** Venn diagram of proteins enriched from the synapse via PD-L1 or PD-1 targeted labeling. PD-L1 and PD-1 hits were determined by a log_2_FC cutoff of 1.32 and 0.58, respectively and p-value <0.05. **b)** GO analysis for enriched biological processes from PD-1 and PD-L1 hits. **c)** Heat map depicting separation scores in primary tumors (TCGA) vs normal tissue (GTEX) for all proximal protein hits when combined with PD-L1. High scores (orange-red) represent strong tumor vs normal separation of the two antigens and are indicative of increased targeted therapy potential. **d)** Single-cell RNA sequencing (scRNAseq) database (Bioturing, Inc) analysis of cell-type expression for genes corresponding to PD-L1 and PD-1 immune synapse proximal proteins. Circle size represents the percentage of a cell type expressing a specified gene. Relative gene expression levels (average rank) are represented by a color scale, yellow (lower) to red (higher).

We next explored gene co-expression of the combined enriched protein hits from **Figure 4a** with PD-L1 in tumor compared to healthy tissue using data from The Cancer Genome Atlas (TCGA) and Genotype-Tissue Expression (GTEX) (**Figure 4c**). Specifically, this was evaluated using a separation score adapted from Dannenfelser et al^40^ (see Methods section). Briefly, the separation score evaluates the tumor-specific targeting potential for two antigens using modalities that allow for Boolean gate antigen binding approaches, e.g. CAR-T, ADC, bispecifics. Specifically, in our case we are evaluating pairs of Antigen 1 “AND” Antigen 2, Antigen 1 “AND NOT” Antigen 2, or Antigen 2 “AND NOT” Antigen 1, where highest scoring pairs represent a good segregation of expression of tumor from normal samples when analyzed in unified expression space. Using this analysis, we observed that POSTN (pancreas), DSG2 (esophagus and lung), PECAM1 (lung), and APOA1 (testis) displayed increased segregation with PD-L1 in tumor types relative to healthy tissue (**Figure 4c**). Furthermore, when we monitored gene expression across all enriched synaptic protein hits, pancreatic cancer stood out with strong gene expression correlation broadly observed in tumor versus healthy tissue compared to other tumor types such as breast cancer (**Supplementary Figure 12**). This result highlights potential novel therapeutic targeting approaches for pancreatic cancer, a tumor type with clinically unmet needs. In addition to co-expression analysis in tumor versus normal tissue, we explored the gene expression profiles of the enriched proteins list across various cell types using a single-cell RNA sequencing (scRNAseq) database to understand gene expression beyond the T and B cell co-culture system used in this study (**Figure 4d** and **Supplementary Table 11**). While expression of PDCD1 and other genes such as CD2, CD8a and ITGAL are specific to lymphoid and T cell lineages, many of the hits such as CD44, CD46, CD47, CD59, BST2 are broadly expressed across different immune cell types.

## Conclusion

In this study we report the development of a flavin-based photoredox labeling platform for proteomic analysis of protein microenvironments within cellular synaptic regions. While most intercellular labeling technologies rely on the use of enzyme-based methods with a primary focus on imaging analysis of physical cell interactions^41–45^, only two enzyme-based approaches (BioID and HRP) have reported proteomic profiling within cell-cell contact environments^11,12^. Whereas those two methods utilized same-cell synapse forming monoculture systems, our approach enriched protein microenvironments from co-culture systems consisting of synapses formed between two different cell types. Furthermore, in contrast to the enzyme-based methods, a key feature of our technology includes the use of visible light for photocatalyst activation. As a result, light can be considered as a discretionary reagent that can be added or removed to instantaneously start or stop the protein labeling reaction and offers the unique ability for temporal control of protein labeling at any given phase of a biological event. Recently, Müller et al.^14^ reported the use of an MHC peptide antigen modified on the N-terminus with a fluorescent dye oxygen photosensitizer for proximal oxidation, followed by biotinylation capture of oxidized protein species in a co-culture system. However, sensitivity of other surface biomolecules such as lipids to singlet oxygen-mediated oxidation^46^, absence of orthogonal selective synaptic labeling evidence (i.e. confocal imaging analysis), and highly variable binding properties displayed by N-terminally modified MHC class I peptides^47^, limits the synaptic labeling selectivity and generalizability of this method. Through our method, we achieve selective and direct synaptic protein labeling on very short timescales (2 minutes) through use of non-blocking VHH modalities that act as spectators on the PD-1 on PD-L1 protein. Moreover, the ability to label from both sides of the synapse with our technology allows for one to adopt this labeling strategy to profile any intercellular junction in a targeting agnostic manner.

While this technology holds promise for unbiased assessment of cis and trans protein interaction environments, disclosure of the initial version of this technology comes with the following considerations.

Our technology requires the use of a non-blocking targeting modality consisting of a VHH-Fc fusion that is site selectively labeled with photocatalyst. As demonstrated with the blocking versus non-blocking VHH-Fc, protein enrichment within the protein microenvironment is sensitive to the nature of the targeting modality. Thus, care must be taken in the selection of targeting modalities that do not interfere with the biology under investigation. Moreover, the small size of the photocatalyst (< 1 kDa) opens up the possibility for direct attachment to other targeting modalities including proteins expressed on the cell surface. Finally, use of the iridium photocatalyst/diazirine biotin probe pair within the immune synapse led to minimal enrichment of surface proteins compared to the RFT system highlighting that while the iridium system is more suitable for target deconvolution or surveying highly localized protein microenvironments, the RFT photocatalyst/biotin phenol probe pair has optimal properties for broad capture of intercellular environments.

Using this flavin-based photocatalytic labeling system, we were able to selectively label, enrich, and identify transcellular protein interactions. This included our intended targeted interaction of PD-L1/PD-1, other known synaptic receptor-ligand pairs and surface proteins, as well as previously unidentified synaptic surface proteins. Ultimately, we envision that the synapse protein microenvironment information generated by this technology within immune synapse and other critical cell-cell contact regions can be combined with genomic and transcriptomic datasets/technologies for a more comprehensive understanding of cell-cell interaction biology to drive novel therapeutic development strategies.

## Acknowledgements

We are very grateful to Nick Haining, Scott Lesley, Erik Hett, and Daria Hazuda (Merck & Co., Inc.) for helpful discussions and Eddie Bowman, Omar Nourzaie, and Sabine Le Saux (Merck & Co., Inc.) for providing protein and cell line reagents.

## Author contributions

O.O.F. and R.C.O. conceived of the work. T.J.B., J.M.B., A.K.O., M.D., C.H.W., T.R.R., O.O.F., and R.C.O. designed and executed experiments. T.J.B., O.O.F., and R.C.O. wrote the manuscript with input from all authors.

## Data Availability

All supplementary figures, experimental details, and synthesis information are included in Supporting Information.

## References

1. Almen, M.S., Nordstrom, K.J., Fredriksson, R. & Schioth, H.B. Mapping the human membrane proteome: a majority of the human membrane proteins can be classified according to function and evolutionary origin. BMC Biol 7, 50 (2009).

2. Dustin, M.L. The immunological synapse. Cancer Immunol Res 2, 1023–33 (2014).

3. Dustin, M.L., Chakraborty, A.K. & Shaw, A.S. Understanding the Structure and Function of the Immunological Synapse. Cold Spring Harbor Perspectives in Biology 2(2010).

4. Alsaab, H.O. et al. PD-1 and PD-L1 Checkpoint Signaling Inhibition for Cancer Immunotherapy: Mechanism, Combinations, and Clinical Outcome. Frontiers in Pharmacology 8(2017).

5. Roux, K.J., Kim, D.I., Raida, M. & Burke, B. A promiscuous biotin ligase fusion protein identifies proximal and interacting proteins in mammalian cells. Journal of Cell Biology 196, 801–810 (2012).

6. Rhee, H.W. et al. Proteomic Mapping of Mitochondria in Living Cells via Spatially Restricted Enzymatic Tagging. Science 339, 1328–1331 (2013).

7. Branon, T.C. et al. Efficient proximity labeling in living cells and organisms with TurboID. Nat Biotechnol 36, 880–887 (2018).

8. Gingras, A.C., Abe, K.T. & Raught, B. Getting to know the neighborhood: using proximity-dependent biotinylation to characterize protein complexes and map organelles. Current Opinion in Chemical Biology 48, 44–54 (2019).

9. Go, C.D. et al. A proximity-dependent biotinylation map of a human cell. Nature 595, 120-+ (2021).

10. Qin, W., Cho, K.F., Cavanagh, P.E. & Ting, A.Y. Deciphering molecular interactions by proximity labeling. Nature Methods 18, 133–143 (2021).

11. Shafraz, O., Xie, B., Yamada, S. & Sivasankar, S. Mapping transmembrane binding partners for E-cadherin ectodomains. Proc Natl Acad Sci U S A 117, 31157–31165 (2020).

12. Loh, K.H. et al. Proteomic Analysis of Unbounded Cellular Compartments: Synaptic Clefts. Cell 166, 1295-+ (2016).

13. Tamura, T., Takato, M., Shiono, K. & Hamachi, I. Development of a Photoactivatable Proximity Labeling Method for the Identification of Nuclear Proteins. Chemistry Letters 49, 145–148 (2020).

14. Muller, M. et al. Light-mediated discovery of surfaceome nanoscale organization and intercellular receptor interaction networks. Nature Communications 12(2021).

15. To, T.L. et al. Photoactivatable protein labeling by singlet oxygen mediated reactions. Bioorganic & Medicinal Chemistry Letters 26, 3359–3363 (2016).

16. Nakane, K. et al. Proximity Histidine Labeling by Umpolung Strategy Using Singlet Oxygen. Journal of the American Chemical Society 143, 7726–7731 (2021).

17. Bechtel, T.J., Reyes-Robles, T., Fadeyi, O.O. & Oslund, R.C. Strategies for monitoring cell-cell interactions. Nature Chemical Biology 17, 641–652 (2021).

18. Ryu, K.A., Kaszuba, C.M., Bissonnette, N.B., Oslund, R.C. & Fadeyi, O.O. Interrogating biological systems using visible-light-powered catalysis. Nature Reviews Chemistry 5, 322–337 (2021).

19. Seath, C.P., Trowbridge, A.D., Muir, T.W. & MacMillan, D.W.C. Reactive intermediates for interactome mapping. Chemical Society Reviews 50, 2911–2926 (2021).

20. Geri, J.B. et al. Microenvironment mapping via Dexter energy transfer on immune cells. Science 367, 1091–1097 (2020).

21. Hope, T. et al. Mechanistic Evidence for a Radical-Radical Recombination Pathway of Flavin-based Photocatalytic Tyrosine Labeling. (American Chemical Society (ACS), 2022).

22. Tay, N. et al. Targeted Activation in Localized Protein Environments via Deep Red Photoredox Catalysis. (American Chemical Society (ACS), 2021).

23. Liu, Z. et al. Spatially resolved cell tagging and surfaceome labeling via targeted photocatalytic decaging. Chem (2022).

24. Oslund, R.C. et al. Detection of cell-cell interactions via photocatalytic cell tagging. Nat Chem Biol 18, 850–858 (2022).

25. Zhao, Y. et al. PD-L1:CD80 Cis-Heterodimer Triggers the Co-stimulatory Receptor CD28 While Repressing the Inhibitory PD-1 and CTLA-4 Pathways. Immunity 51, 1059–1073 e9 (2019).

26. Orme, J.J. et al. ADAM10 and ADAM17 cleave PD-L1 to mediate PD-(L)1 inhibitor resistance. Oncoimmunology 9(2020).

27. Verschueren, E. et al. The Immunoglobulin Superfamily Receptome Defines Cancer-Relevant Networks Associated with Clinical Outcome. Cell 182, 329-+ (2020).

28. Platt, J.L. et al. C3d regulates immune checkpoint blockade and enhances antitumor immunity. JCI Insight 2(2017).

29. Aust, S. et al. Absence of PD-L1 on tumor cells is associated with reduced MHC I expression and PD-L1 expression increases in recurrent serous ovarian cancer. Scientific Reports 7(2017).

30. Roemer, M.G.M. et al. Major Histocompatibility Complex Class II and Programmed Death Ligand 1 Expression Predict Outcome After Programmed Death 1 Blockade in Classic Hodgkin Lymphoma. Journal of Clinical Oncology 36, 942-+ (2018).

31. Grande, F. et al. CCR5/CXCR4 Dual Antagonism for the Improvement of HIV Infection Therapy. Molecules 24(2019).

32. Tamagnone, L. & Franzolin, G. Targeting Semaphorin 4D in Cancer: A Look from Different Perspectives. Cancer Research 79, 5146–5148 (2019).

33. Barlesi, F. et al. 291 Phase Ib study of selicrelumab (CD40 agonist) in combination with atezolizumab (anti-PD-L1) in patients with advanced solid tumors. Journal for ImmunoTherapy of Cancer 8, A178–A178 (2020).

34. Ma, H.S. et al. A CD40 Agonist and PD-1 Antagonist Antibody Reprogram the Microenvironment of Nonimmunogenic Tumors to Allow T-cell-Mediated Anticancer Activity. Cancer Immunology Research 7, 428–442 (2019).

35. UniProt, C. UniProt: the universal protein knowledgebase in 2021. Nucleic Acids Res 49, D480–D489 (2021).

36. Bausch-Fluck, D. et al. The in silico human surfaceome. Proc Natl Acad Sci U S A 115, E10988–E10997 (2018).

37. Borst, J. et al. Complexity of T-Cell Receptor Recognition Sites for Defined Alloantigens. Journal of Immunology 139, 1952–1959 (1987).

38. Szklarczyk, D. et al. STRING v11: protein-protein association networks with increased coverage, supporting functional discovery in genome-wide experimental datasets. Nucleic Acids Res 47, D607–d613 (2019).

39. Gene Ontology, C. The Gene Ontology resource: enriching a GOld mine. Nucleic Acids Res 49, D325–D334 (2021).

40. Dannenfelser, R. et al. Discriminatory Power of Combinatorial Antigen Recognition in Cancer T Cell Therapies. Cell Systems 11, 215-+ (2020).

41. Cho, K.F. et al. Split-TurboID enables contact-dependent proximity labeling in cells. Proceedings of the National Academy of Sciences of the United States of America 117, 12143–12154 (2020).

42. Liu, Q. et al. A proximity-tagging system to identify membrane protein-protein interactions. Nature Methods 15, 715-+ (2018).

43. Martell, J.D. et al. A split horseradish peroxidase for the detection of intercellular protein-protein interactions and sensitive visualization of synapses. Nat Biotechnol 34, 774–80 (2016).

44. Liu, D.S., Loh, K.H., Lam, S.S., White, K.A. & Ting, A.Y. Imaging trans-cellular neurexin-neuroligin interactions by enzymatic probe ligation. PLoS One 8, e52823 (2013).

45. Carpenter, M.A. et al. Protein Proximity Observed Using Fluorogen Activating Protein and Dye Activated by Proximal Anchoring (FAP–DAPA) System. ACS Chemical Biology (2020).

46. Di Mascio, P. et al. Singlet Molecular Oxygen Reactions with Nucleic Acids, Lipids, and Proteins. Chemical Reviews 119, 2043–2086 (2019).

47. Garstka, M.A. et al. The first step of peptide selection in antigen presentation by MHC class I molecules. Proceedings of the National Academy of Sciences of the United States of America 112, 1505–1510 (2015).

