## Supplementary Information for "Proteomic Mapping of Intercellular Synaptic Environments via Flavin-Dependent Photoredox Catalysis"

### Supplementary Figures

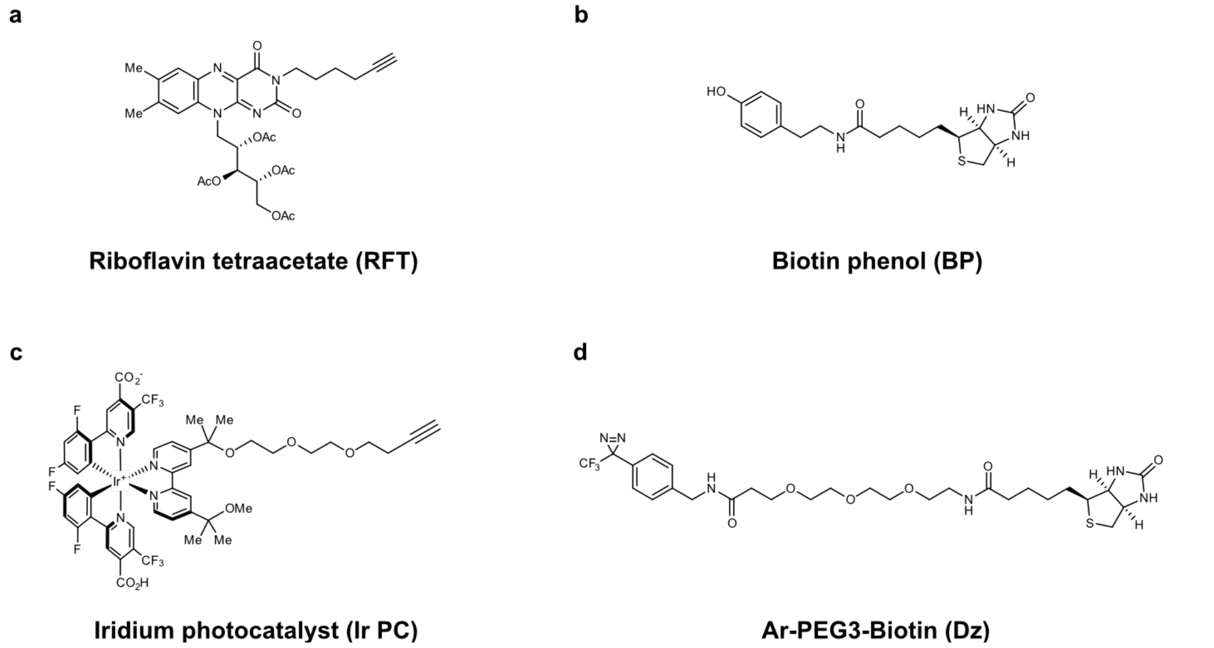

**Supplementary Figure 1. Chemical structures of photocatalysts and small molecule probes. a)** Riboflavin tetraacetate (RFT, photocatalyst used in PhoTag). **b)** Biotin phenol (BP, biotin containing probe used in PhoTag). **c)** Iridium photocatalyst (Ir PC, used with μMap). **d)** Ar-PEG3-Biotin (Dz, a biotin containing probe used with μMap).

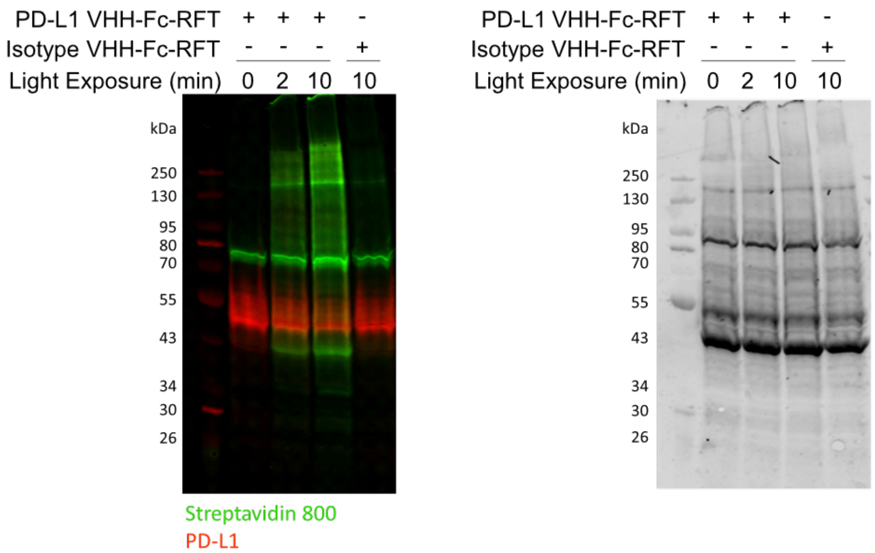

**Supplementary Figure 2. Western blot analysis (left panel) with total protein stain (right panel).**

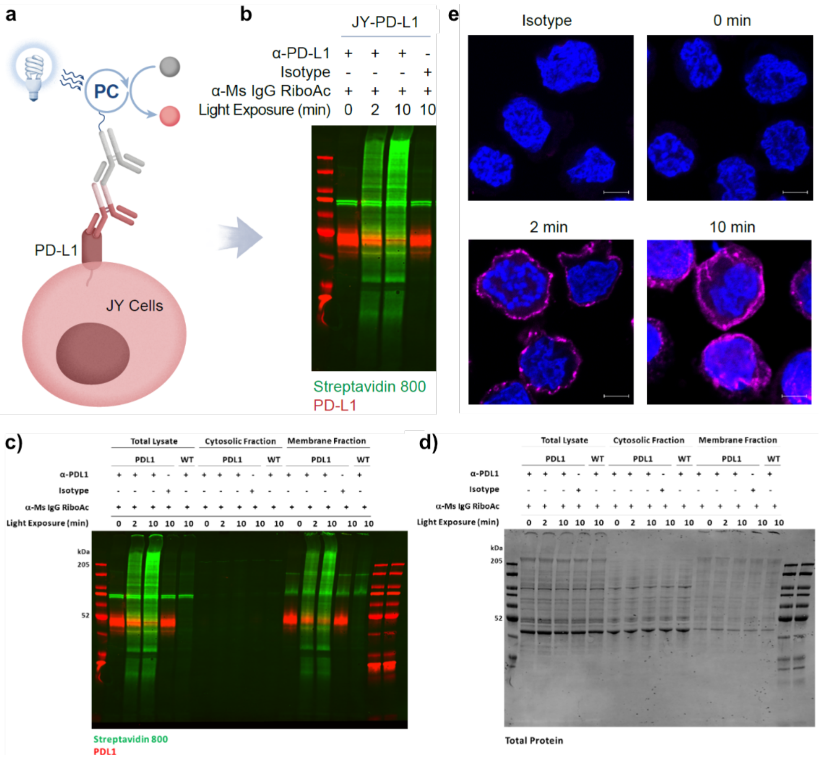

**Supplementary Figure 3. PD-L1 targeted photocatalytic proximity labeling on the surface of JY B cells using a two-antibody system. a)** Schematic depicting the targeted labeling of PD-L1 proximal proteins on the surface of JY PD-L1 cells using a PD-1 blocking α-PD-L1 primary antibody and goat α-mouse secondary antibody conjugated to riboflavin tetraacetate (RFT). After antibody-based targeting, biotin phenol is added, and the cells are irradiated with blue light to induce photocatalytic proximity labeling events. **b)** Western blot analysis of PD-L1 targeted labeling on JY B cells. Protein biotinylation (green) increases as a function of blue-light irradiation time in PD-L1 targeted samples. Minimal background labeling is observed with the use of a non-binding isotype control. **c)** In addition to total lysates, cytosolic and membrane fractions were isolated to determine the subcellular selectivity of these biotinylation events. After photolabeling, total cell lysate, cytosolic lysate, or enriched membrane lysate was generated and analyzed by western blot for biotinylation and PD-L1. The absence of biotinylation signal in the cytosolic fraction supports primary biotinylation at the membrane surface. **d)** Total protein stain of panel c. **e)** Confocal microscopy imaging of PD-L1 targeted labeling on JY cells reveals that biotinylation events (magenta stain) are confined to the cell surface. Nuclei are labeled with Hoechst stain (blue) and scale bars indicate 5 µm.

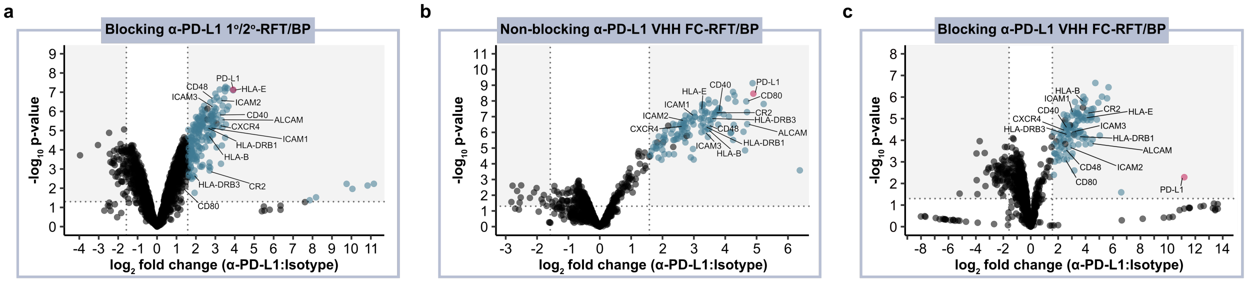

**Supplementary Figure 4. Quantitative proteomic analysis of enriched proteins from PD-L1 targeting on JY PD-L1 cells with various targeting modalities. a)** JY PD-L1 cells targeted with a PD-L1 blocking α-PD-L1 primary antibody or isotype control and goat α-mouse-RFT secondary antibody. **b)** JY PD-L1 cells targeted with a non-blocking α-PD-L1 VHH-Fc-RFT or isotype control. **c)** Blocking α-PD-L1 VHH-Fc-RFT data (**from Figure 2b**) was included to compare each PD-L1 targeting modality. **a-c**) Samples were irradiated with blue light for 2 minutes in the presence of biotin phenol. Biotinylated proteins were enriched by streptavidin beads and subjected to tandem mass tag (TMT)-based quantitative proteomic analysis. Protein enrichment is visualized using a volcano plot of statistical significance vs. fold enrichment (α-PD-L1:Isotype). Significantly enriched cell-surface proteins ($p$-value < 0.05 and $\log_2 FC > 1.58$) are indicated as blue dots and PD-L1 is indicated as a red dot (n = 3 experiments). Protein localization to the cell surface was determined by Uniprot[1] and the Surfaceome[2].

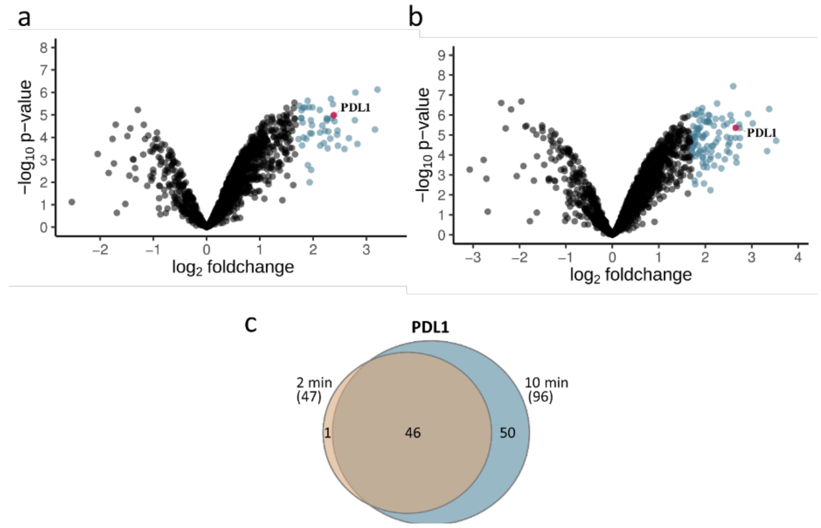

**Supplementary Figure 5. Quantitative proteomic analysis of enriched proteins from PD-L1 targeting on JY PD-L1 cells with 2 min vs 10 min irradiation.** In a separate experiment, we explored the time-dependent biotinylation of proteins proximal to PD-L1 via MS-based proteomics. JY PD-L1 cells were targeted with α-PD-L1 primary antibody or isotype control and goat α-mouse-RFT secondary antibody. Following the addition of biotin phenol and visible light irradiation for 2 or 10 min, biotinylated proteins were subjected to streptavidin-bead enrichment, followed by quantitative TMT-based proteomic analysis. Volcano plots of significance vs. fold-enrichment were generated to visualize protein enrichment from **a)** 2 min or **b)** 10 min of light-induced labeling. Significantly enriched proteins (defined as having an FDR corrected p-value < 0.05, displaying enrichment of > 2.5-fold above controls, n = 3 experiments) are indicated in blue and PD-L1 in red. The number of significantly enriched proteins detected in the 10-min labeling experiment is larger than that of the 2-min experiment. **c)** Venn diagram displaying the overlap between significantly enriched protein populations at 2-min and 10-min of labeling, showing that the larger set of enriched proteins found following 10-min of labeling contains the vast majority of the smaller 2-min labeling subset. Significantly enriched protein population sizes are indicated numerically on the diagram.

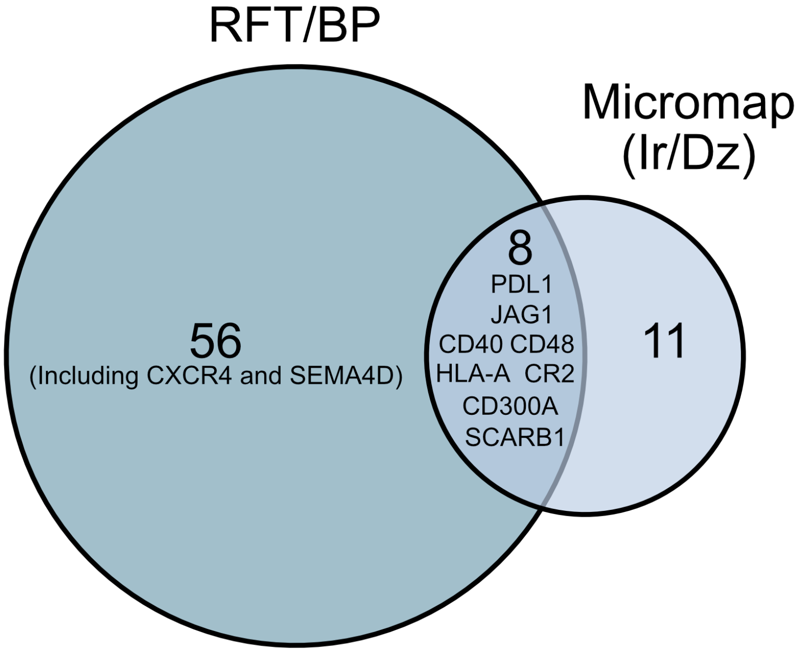

**Supplementary Figure 6. Comparison of PD-L1 proximal proteins identified by photoproximity labeling technologies on JY PD-L1 cells.** The identity of proteins sharing statistically significant enrichment across all three α-PD-L1 targeting modalities (blocking two-antibody, blocking VHH-Fc, and non-blocking VHH-Fc) with the RFT/BP proximity labeling approach were compared with significantly enriched proteins identified by an alternative photocatalytic proximity labeling strategy that used an α-PD-L1 two-antibody targeting modality and iridium photocatalyst/biotin diazirine (Ir/Dz) probe system[3]. While both approaches enriched PD-L1 and a subset of surface proteins (including CD40), the RFT/BP-based proximity labeling resulted in a larger set of statistically significantly enriched proteins including CXCR4 and SEMA4D. Significantly enriched protein population sizes are indicated numerically on the diagram.

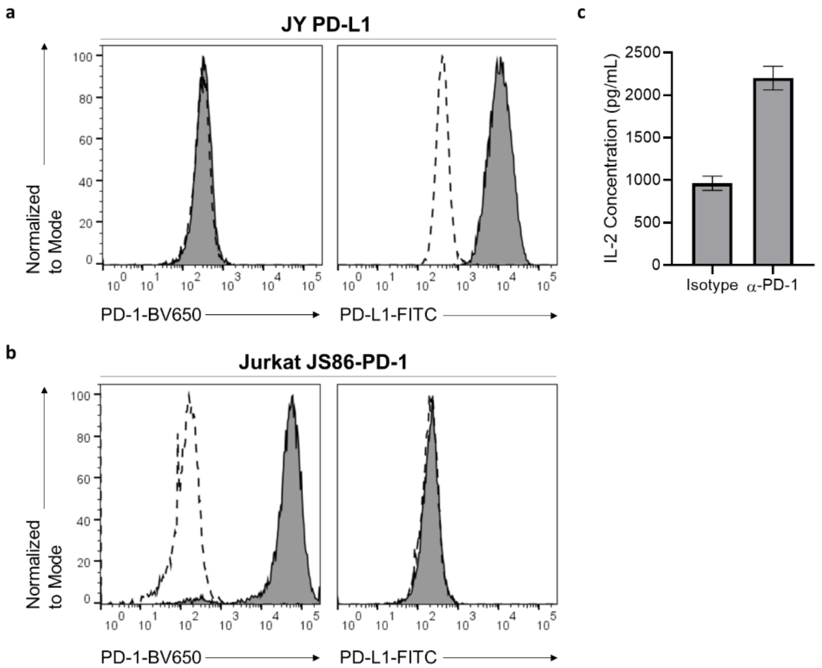

**Supplementary Figure 7. Characterization of Jurkat JS86-PD-1 and JY PD-L1 co-culture system. a)** Surface PD-1 and PD-L1 expression on JY PD-L1 cells was measured by flow cytometry. High expression of PD-L1 was detected compared to Isotype control or α-PD-1 antibody. **b)** Surface PD-1 and PD-L1 expression on Jurkat JS86-PD-1 cells was measured by flow cytometry. High expression of PD-1 was detected compared to Isotype control or α-PD-L1 antibody. **c)** Jurkat JS86-PD-1 cells were co-incubated with JY PD-L1 cells for ~24 hours followed by detection of IL-2 production using a human IL-2 ELISA kit. Bar plot analysis shows increased IL-2 expression with an α-PD-1 antibody compared to Isotype treatment demonstrating that PD-1/PD-L1 signaling is functionally intact. Error bars represent standard deviation of n = 3 experiments.

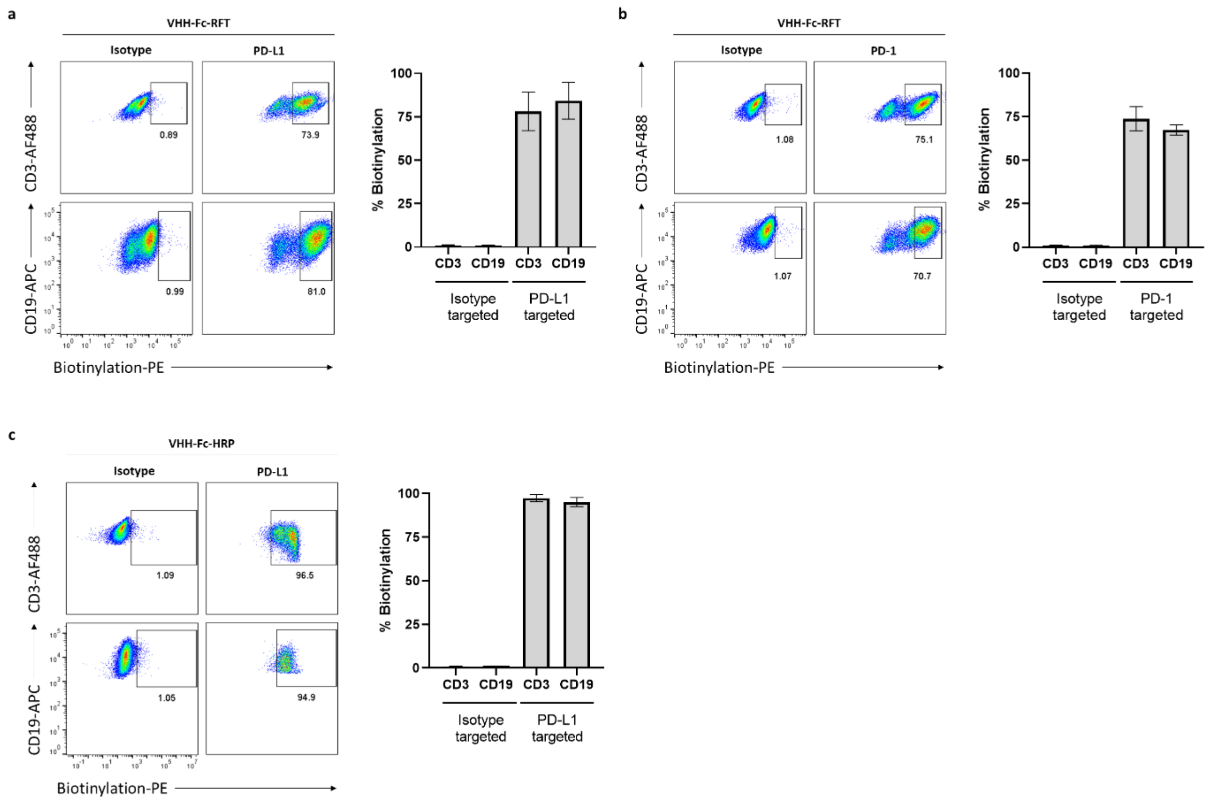

**Supplementary Figure 8. Flow cytometry analysis of *cis* and *trans* labeling of JY PD-L1 and Jurkat JS86-PD-1 cells with non-blocking α-PD-L1 VHH-Fc-RFT or VHH-Fc-HRP.** a) JY PD-L1 cells ($CD19^+$) were pre-labeled with isotype control (Isotype VHH-Fc-RFT) or α-PD-L1 VHH-Fc-RFT, followed by 30-minute co-culture with Jurkat JS86-PD-1 cells ($CD3^+$) and visible light labeling for 2 minutes. b) Jurkat JS86-PD-1 cells ($CD3^+$) were pre-labeled with isotype control (Isotype VHH-Fc-RFT) or α-PD-1 VHH-Fc-RFT, followed by 30-minute co-culture with JY PD-L1 cells ($CD19^+$) and visible light labeling for 2 minutes. c) JY PD-L1 cells ($CD19^+$) were pre-labeled with isotype control (Isotype VHH-Fc-HRP) or α-PD-L1 VHH-Fc-HRP, followed by 30-minute co-culture with Jurkat JS86-PD-1 cells ($CD3^+$) and labeling for 1 minute. *cis* and *trans* labeling of JY PD-L1 and Jurkat JS86-PD-1 cells was observed when targeting either PD-1 or PD-L1 and or either catalyst, RFT or HRP. In each panel, flow cytometry dot plots are shown for each condition along with bar plots of triplicate analysis (error bars represent standard deviation).

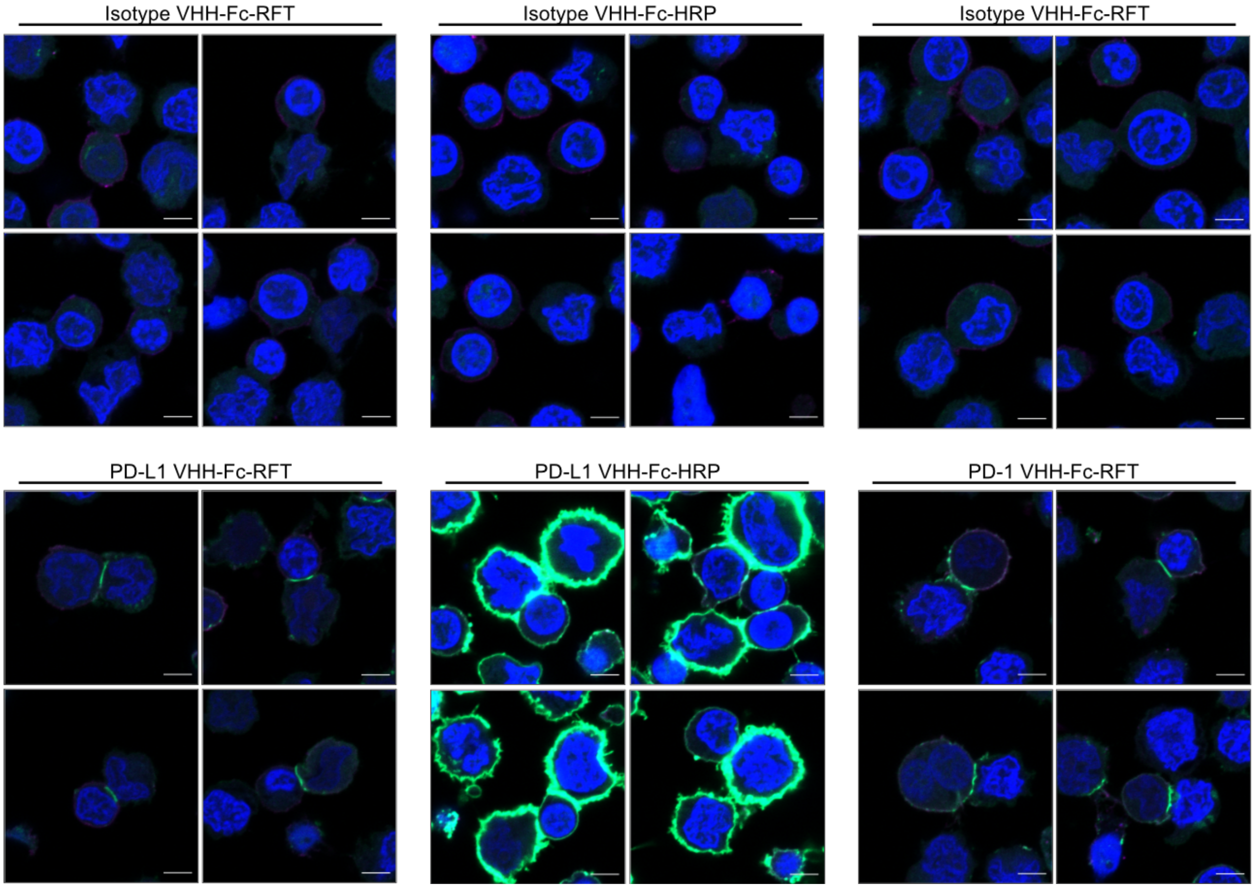

**Supplementary Figure 9. Confocal microscopy analysis of *cis* and *trans* labeling of JY PD-L1 and Jurkat JS86-PD-1 cells with non-blocking α-PD-L1 VHH-Fc-RFT or VHH-Fc-HRP.** JY PD-L1 cells were targeted with Isotype VHH-Fc-RFT or VHH-Fc-HRP, non-blocking α-PD-L1 VHH-Fc-RFT or VHH-Fc-HRP while Jurkat JS86-PD-1 cells were targeted with Isotype VHH-Fc-RFT or non-blocking α-PD-1 VHH-Fc-RFT. Labeled JY PD-L1 cells and Jurkat JS86-PD-1 cells were incubated with non-labeled Jurkat JS86-PD-1 cells and JY PD-L1 cells, respectively, for 30 min and subjected to proximity labeling workflows. Cells were irradiated with blue light for 2 min. Biotinylation events (green) were visualized by confocal microscopy. Immune synapse-selective biotinylation events were observed when using α-PD-L1 or α-PD-1 VHH-Fc-RFT modalities. Excessive biotinylation beyond the immune synapse was found when using α-PD-L1 VHH-Fc-HRP. Nuclei are labeled with Hoechst stain (blue), CD3 on T cells were also stained (purple), and scale bars indicate 5 µm. Four different images are shown for each condition.

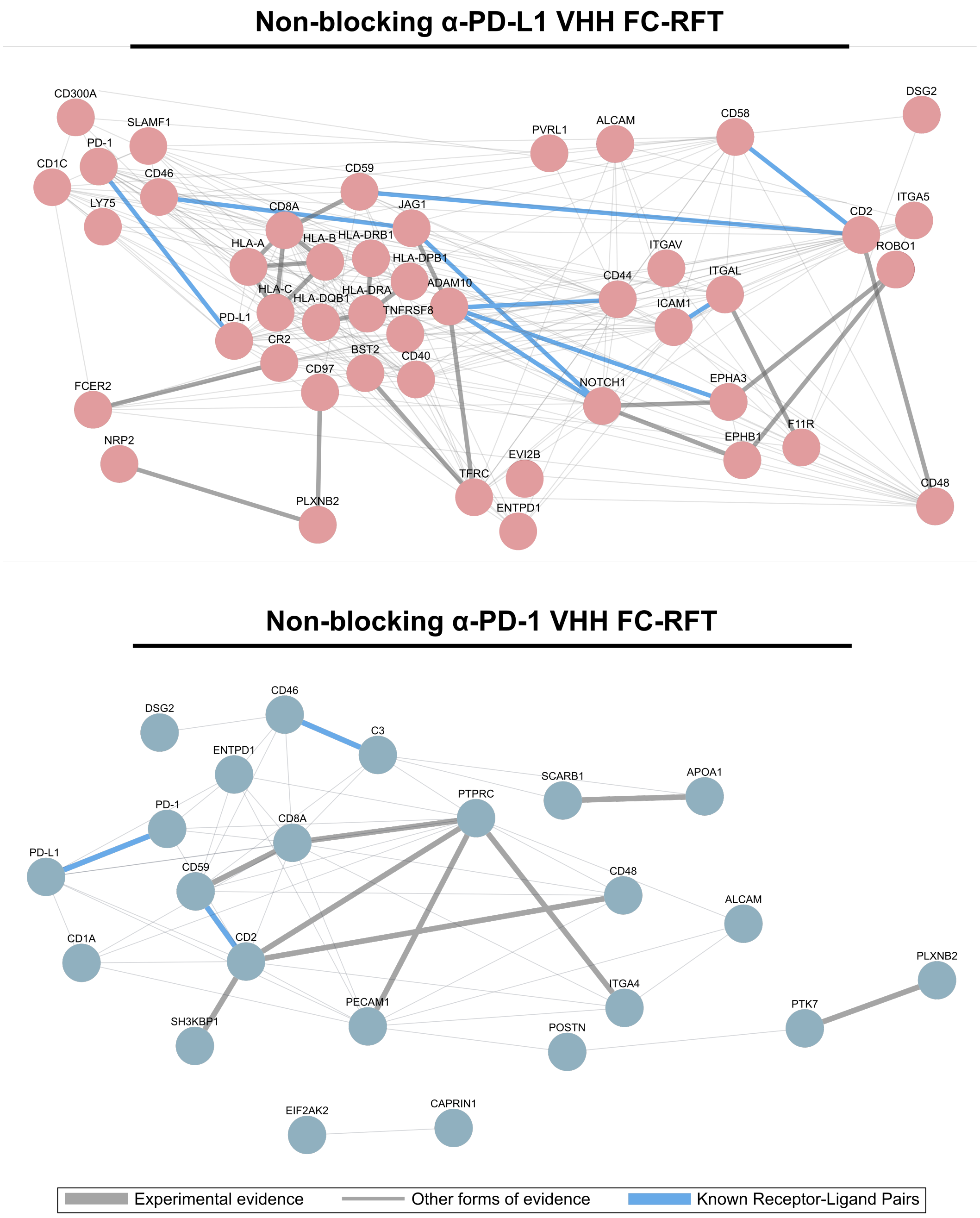

**Supplementary Figure 10. Protein-protein interaction analysis of enriched proteins using PD-L1 or PD-1 targeted photocatalytic proximity labeling within JY PD-L1-Jurkat JS86-PD-1 immune synapses.** STRING analysis of *cis* and *trans* protein interactions were plotted for intercellular PD-L1 targeted labeling (top) or intercellular PD-1 targeted labeling (bottom). Only enriched proteins with at least one interaction were included. Thick edges denote protein interactions with supporting experimental evidence. Thin lines represent interactions based on all other sources of evidence as indicated by the STRING database[4]. Known intercellular receptor-ligand pairs as determined by CellTalkDB[5] are further indicated with a blue line.

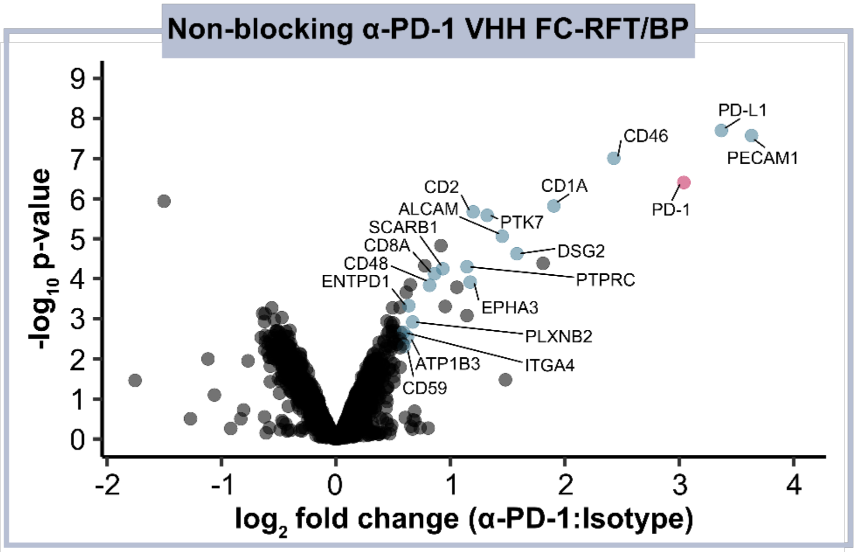

**Supplementary Figure 11. Photocatalytic proximity labeling of PD-1 microenvironments within the immunological synapse of Jurkat JS86-PD-1 T cells and JY PD-L1 B cells.** Jurkat JS86-PD-1 cells pre-treated with non-blocking α-PD-1 VHH-Fc-RFT were co-incubated with JY PD-L1 cells for 30 min. Following incubation, cells were treated with biotin phenol and irradiated with blue light for 2 min. Biotinylated proteins were subjected to streptavidin enrichment, followed by TMT-based proteomic analysis. A volcano plot of significance vs. fold-enrichment was generated to visualize protein enrichment. Statistically significantly enriched cell surface proteins (p-value <0.05 and $log_2FC > 0.58$, n = 3 experiments) are indicated in blue and PD-1 in red. Protein localization to the cell surface was determined by Uniprot[1] and the Surfaceome[2].

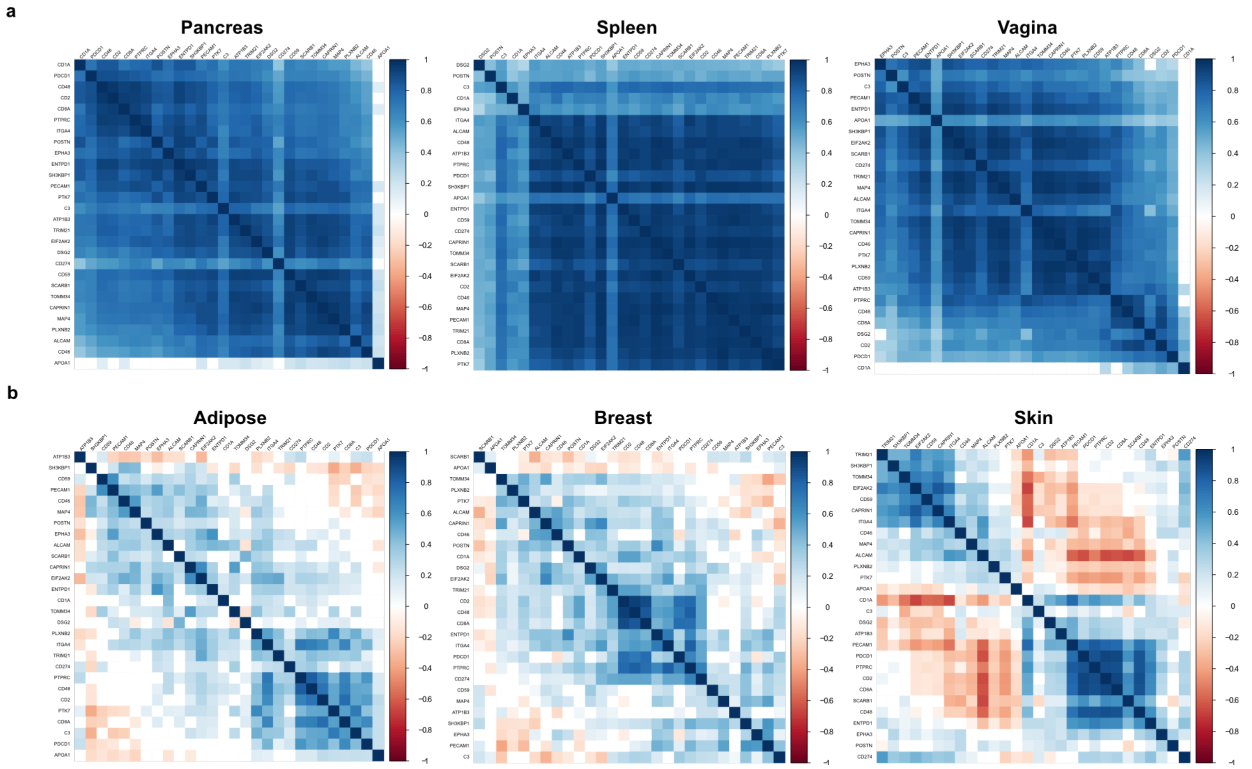

**Supplementary Figure 12. Gene expression correlation analysis of all significantly enriched proteins from PD-1 and PD-L1 targeted photocatalytic proximity labeling within the immune synapse.** Correlations in gene expression between proteins significantly enriched for by PD-1 or PD-L1 targeted labeling in the immune synapse was assessed within various tumor tissue types. Positive or negative correlations are indicated in blue or red, respectively. In addition to pancreatic tumors, global positive correlations in gene expression were also observed within spleen and vaginal tumors. Meanwhile, mixed correlations in gene expression were generally observed across several tumor tissue types such as adipose, breast, and skin tumors.

### Materials and Methods

#### General Materials

Ascorbic acid (BP321-500) and Ethanol (BP2818100) was purchased from Fisher Scientific (Pittsburgh, PA). Eppendorf Protein LoBind tubes (Z666505-100EA), Trolox (238813), Biotin phenol (SML2135), 5M Sodium Chloride (71386), 20% Sodium dodecyl sulfate (SDS, 05030) and Hydrogen peroxide (88597) were purchased from Sigma-Aldrich (St. Louis, MO). Sodium azide (14314) was purchased from Alfa Aesar (Haverhill, MA). RIPA Buffer (89900), Click-iT™ Protein Reaction Buffer Kit (C10276), DPBS (14190144), Pierce™ BCA Protein Assay kit (23227), Zeba Spin desalting column (87769), iBright Prestained Protein Ladder (LC5615) and Dithiothreitol (DTT, 15508013) were purchased from Thermo Scientific (Rockford, IL). TBST (IBB-581X) was purchased from Boston BioProducts (Ashland, MA). Criterion XT 12% Tris TGX precast gels (5671044) and 4x Laemmli sample buffer (161-0747) were purchased from Bio-Rad (Hercules, CA). Jurkat NF-κβ GFP cells and JY PD-L1 cells were a gift from Rene De Waal Malefyt and Sabine Le Saux (MRL, Merck & Co., Inc., Palo Alto, CA, USA).

#### General Cell Culture

JY PD-L1 and JY WT cells were cultured in RPMI 1640 1x with L-glutamine (Corning, Cat: 10-040-CV) supplemented with 10% fetal bovine serum (FBS) (HyClone, Cat: SH30910.03), 100 IU Penicllin/100µg/ml Streptomycin (1% final concentration from 100x stock, Corning, Cat: 30-002-CI), 2mM L-Glutamine (from 200mM stock, Lonza, Cat: 17-605E), 1x MEM Non-Essential Amino Acid solution (MEM NEAA) (from 100x stock, Gibco, Cat: 11140-050) and 1mM sodium pyruvate (from 100mM stock, Gibco, Cat: 11360-070). Jurkat PD-1 cells were cultured in RPMI 1640 1x with L-glutamine (Corning, Cat: 10-040-CV) supplemented with 10% FBS (HyClone, Cat: SH30910.03), 250 µg/ml geneticin (from 50 mg/ml stock, Gibco, Cat: 10131-035) and 250 µg/ml Hygromycin B (from 50 mg/ml stock, Invitrogen, Cat: 10687010). Jurkat NF-κβ GFP cells were cultured in RPMI 1640 1x with L-glutamine (Corning, Cat: 10-040-CV) supplemented with 10% FBS (Hyclone, Cat: SH30910.03) and 100 IU Penicillin/100 µg/ml Streptomycin (1% final concentration from a 100x stock, Corning, Cat: 30-002-CI).

Cell culture media was filter-sterilized via 0.2 µm Nalgene™ Rapid-Flow™ Sterile Disposable Filter Units with PES Membrane, 500ml capacity (Thermo Scientific, Cat: 569-0020) or 1,000ml capacity (Thermo Scientific, Cat: 567-0020).

All cell cultures were grown at 37$^{o}$C with 5% $CO_2$ in 25 cm$^2$ (Corning, Cat: 430639), 75 cm$^2$ (Corning, Cat: 430641U) or 150 cm$^2$ (Corning, Cat: 430825) vented cap sterile cell culture flasks. For passaging, cells were collected and centrifuged at 300g x 5 min. Supernatant was removed and cells were diluted to the desired concentration.

#### Generation of Co-Culture System Using Jurkat JS86-PD-1 and JY PD-L1 cells

The Jurkat cell line expressing the HLA-A02-recognizing receptor, JS86[1], and the CD8 receptor was a gift from Alberto Visintin (Merck & Co., Inc., Boston, MA, USA). These cells were further engineered to overexpress PD-1 and Cas9 via lentiviral delivery with reagents obtained from the Genetic Perturbation Platform at the Broad Institute.

Cas9 was introduced via transduction using 5e5 cells in infection media (1.5ml RPMI+10% FBS supplemented with 5µg/ml polybrene (Sigma, Cat: SIALMSD-TR-1003-G)) and 0.5ml of lentivirus. The next day, the cells were gently centrifuged, and the infection media was replaced with 2ml fresh complete media. The following day, cell media supplemented with hygromycin B (Invitrogen, Cat: 10687010) was added to the cells and passaging was performed as needed for the next 4 days in selection media for a total of 7 days. Functional validation of Cas9 was performed using a CRISPRtest™ Functional Cas9 Activity Kit (Cellecta, Cat: CRTEST) per the manufacturer's instructions. Cas9$^+$ cells were expanded, and PD-1 was introduced via lentivirus as indicated above and selected for geneticin (neomycin) (Invitrogen, Cat: 10131-035) resistance. Single cell sorting of Cas9$^+$ PD-1$^+$ clones was performed using a BD FACSMelody with BD FACSChorus software and PD-1 levels evaluated using a BD FACSCelesta with BD FACSDiva software. Data was analyzed using FlowJo, v10.

##### Cell Surface Staining for PD-L1 or PD-1 on JY PD-L1 or Jurkat JS86-PD-1 Cells

1 million JY PD-L1 or Jurkat JS86-PD-1 cells per sample were centrifuged for 5 min at 500xg, 4°C, washed 1x with 200µl 1x DPBS, and centrifuged again. The cell pellet was resuspended with 100µl of a 1:500 dilution of Zombie Violet Fixable Viability Kit (BioLegend, Cat: 423113) for 15 minutes at room temperature (RT), covered from light. The volume per well was brought up to 200µl with 1x DPBS, centrifuged for 5 min at 500xg, 4°C, washed 1x with 200µl 1x DPBS, and centrifuged again. Each sample was resuspended in 100µl of Fc Block (BD Biosciences, Cat: 564220) at a 1:100 dilution in Stain Buffer (BD Biosciences, Cat: 554656) for 20 minutes at 4°C. The volume per well was brought up to 200µl with Stain Buffer, centrifuged for 5 min at 500xg, 4°C, washed 1x with 200µl Stain Buffer, and centrifuged again. The cells were stained in 100µl Stain Buffer containing a 1:20 dilution of BV650 mouse anti-human PD-1 antibody, clone MIH-4 (BD Biosciences, Cat: 564324) and a 1:20 dilution of FITC mouse anti-human PD-L1 antibody, clone MIH-1 (BD Biosciences, Cat: 558065), or a 1:20 dilution of BV650 Mouse IgG1, k Isotype control antibody (BD Biosciences, Cat: 563231) and a 1:20 dilution of FITC Mouse IgG1, k Isotype control antibody (BD Biosciences, Cat: 555748). Samples were incubated for 30 minutes at 4°C, covered from light.

The volume per well was brought up to 200µl with Stain Buffer, centrifuged for 5 min at 500xg, 4°C, washed 1x with 200µl Stain Buffer, and centrifuged again. The final cell pellets were resuspended in 200µl Stain Buffer and transferred to 5ml FACS tubes (Fisherbrand, Cat: 14-956-3D). Samples were acquired in a BD FACSCelesta with BD FACSDiva software. Data was analyzed using FlowJo, v10.

##### Functional Validation of JY PD-L1 – Jurkat JS86-PD-1 Co-Culture System

Functional validation of the JY PD-L1-Jurkat JS86-PD-1 two-cell system was achieved by monitoring IL-2 production induced by co-incubation of these two cell lines in the presence or absence of PD-1 antibody-mediated blockade. Briefly, 45µl of an 8 million cells/ml stock of Jurkat JS86-PD-1 cells in assay medium (RPMI 1640, Corning, Cat: 10-040-CV + 10% dialyzed FBS, HyClone, Cat: SH30079.03) were mixed with 45µl of mouse IgG1 κ isotype control antibody, clone MOPC-21 (BD Biosciences, Cat: 556648), or α-human PD-1 antibody (Creative Biolabs, Cat: TAB-770) at 10µg/ml in assay medium and incubated for 30 min at room temperature (RT) in a 96-well, U-bottom plate.

After the 30 min incubation, 125µl of JY PD-L1 cells were aliquoted per well in a separate 96-well, U-bottom plate and mixed with 25µl of antibody-loaded Jurkat JS86-PD-1 cells (100,000 cells per cell line). The plate was incubated for ~24 hours at 37°C + 5% $CO_2$. The following day, the plate was centrifuged for 1 min at 1,100 RPM and RT. 50µl of culture supernatant per well were used to measure IL-2 production using a Human IL-2 ELISA kit (Thermo Fisher, Cat: EH2IL25) per manufacturer's instructions. Absorbance at 550 nm was subtracted from measurements at 450 nm and the final values were used as a readout for IL-2 production. Data was analyzed and graphed using MS Excel and GraphPad Prism software (8.1.1).

#### General Synthetic Information

The primary photocatalyst used for photocatalytic proximity labeling, Riboflavin tetraacetate (RFT), was synthesized as previously described.[6] The iridium photocatalyst and biotin-containing diazirine probe (Ar-

PEG3-Biotin) used to compare different proximity labeling technologies within the immune synapse was synthesized as previously described in Geri, J.B. et al.[3]

#### Preparation of secondary antibody-RFT conjugate

300µl of polyclonal goat α-mouse IgG (Millipore, Cat: AP124) was combined with 30µl of 1M sodium bicarbonate buffer (pH 8.5) in a Protein LoBind tube. 3µl of 100mM azidobutyric acid NHS ester (prepared in DMSO) (Enamine, Cat: EN300-265680) was added and the reaction mixture was incubated for 1.5 hours in the dark. After 1.5 hours, an additional 3µl of 100 mM azidobutyric acid NHS ester linker was added, and the sample was incubated for 1.5 hours in the dark. After the second incubation, the sample was buffer exchanged into 50mM Tris pH 8.0 using a Zeba Spin desalting column (Thermo Scientific, Cat: 87769, 2ml column, 40,000 MWCO), according to the manufacturer's instructions. 200µl of the antibody-azide conjugate was transferred to a Protein LoBind tube for click chemistry azide-alkyne cycloaddition of RFT using the Click-iT™ Protein Reaction Buffer Kit (Invitrogen, Cat: C10276). Briefly, 300 µM RFT (final concentration from 5 mM stock in DMSO) was added to the antibody-azide conjugate and vortexed. 12.5µl of copper sulfate was added and the sample was vortexed. 12.5µl of additive 1 was added, the sample was vortexed, and incubated for 3 min at RT. Following incubation, 15µl of additive 2 was added to the reaction mixture, the sample was vortexed, and incubated in the dark for 30 min at RT. After incubation, the sample was buffer exchanged into 1x DPBS using a Zeba Spin desalting column (2ml, 40,000 MWCO) according to the manufacturer's instructions. The final protein concentration of the secondary antibody-RFT conjugate was determined using the Pierce™ BCA Protein Assay kit (Thermo Scientific, Cat: 23227) according to the manufacturer's instructions. RFT photocatalyst concentration was determined by measuring the absorbance at 450nm (A450) and compared to a standard curve consisting of known free RFT concentrations. The micromolar concentration of RFT was divided by the micromolar concentration of antibody to determine the antibody:RFT ratio. We routinely obtained an antibody:RFT ratio of ~1:5.

#### Preparation of VHH-Fc-RFT conjugates

RFT DBCO was site selectively appended to Isotype VHH-Fc-$N_3$[6], non-blocking α-PD-L1 VHH-FC-$N_3$[6], blocking α-PD-L1 VHH-FC-$N_3$[6], or non-blocking α-PD-1 VHH-FC-$N_3$[6] as previously described[6]. Briefly, 200µl of VHH-Fc-$N_3$ conjugate (1 mg/ml in 1x DPBS) was combined with 600µM RFT DBCO and incubated overnight at 4$^o$C. The labeled product, VHH-Fc-RFT, was buffer exchanged into DPBS using a Zeba Spin desalting column (2ml, 40,000 MWCO) according to the manufacturer's instructions. VHH-Fc concentration was determined using the Pierce™ BCA Protein Assay kit according to the manufacturer's instructions, and RFT concentration was determined by measuring absorbance at 450nm (A450) and comparing it to a standard curve consisting of known free RFT concentrations. The micromolar concentration of RFT was divided by the micromolar concentration of VHH-Fc to determine the VHH-Fc:RFT ratio. A VHH-Fc:RFT ratio of 1:3 for each type of VHH-Fc-RFT conjugate was routinely obtained.

#### Preparation of VHH-Fc-Ir conjugate

The iridium photocatalyst, Ir PC alkyne, was site selectively conjugated to non-blocking α-PD-L1 VHH-Fc-$N_3$[6] or Isotype VHH-Fc-$N_3$[6] via click chemistry azide-alkyne cycloaddition with the Click-iT™ Protein Reaction Buffer Kit according to the manufacturer's instructions. Briefly, 200µl of VHH-Fc-$N_3$ was combined with 300 µM Ir PC alkyne in a Protein LoBind tube and vortexed. Next, 12.5µl of copper sulfate and 12.5µl of additive 1 were added to the mixture with the sample being vortexed after the addition of each reagent. After adding additive 1, the sample was incubated for 3 min followed by the addition of 15µl of additive. The sample was vortexed and incubated for 30 min in the dark at RT. The sample was then buffer exchanged into 1x DPBS using a Zeba Spin desalting column (2ml, 40,000 MWCO) according to the manufacturer's instructions. VHH-Fc concentration was determined using the Pierce™ BCA Protein Assay kit according to the manufacturer's instructions, and Ir PC concentration was determined by measuring absorbance at 350nm (A350) and comparing it to a standard curve consisting of known free Ir PC concentrations. The micromolar concentration of Ir PC alkyne was divided by the micromolar concentration of VHH-Fc to determine the VHH-Fc:Ir PC ratio. We routinely obtained a VHH-Fc:Ir PC ratio of 1:3 for each type of VHH-Fc-Ir conjugate.

#### Preparation of VHH-Fc-HRP conjugate

Non-blocking α-PD-L1 VHH-Fc-$N_3$[6] or Isotype VHH-Fc-$N_3$[6] was conjugated to horseradish peroxidase (HRP) using an HRP Conjugation Kit (Abcam: ab102890) according to the manufacturer's instructions as previously disclosed[6]. Briefly 10µl of modifier reagent was combined with 100µl of VHH-Fc-$N_3$ (1 mg/ml in DPBS). This solution was then mixed with 100 µg of HRP and incubated at RT overnight in the dark. Following incubation, 10µl of quencher solution was added and the sample was incubated for 30 min. VHH-Fc-HRP was stored at 4$^{o}$C until use.

#### Western Blotting

Samples were loaded onto Criterion TGX 12% gels (Bio-Rad, Cat: 5671044) and electrophoresed for 60 minutes at 180V. The proteins were blotted onto a PVDF membrane (Thermo Fisher, Cat: IB24001) and the membranes were blocked with 50ml of Tris Buffered Saline-Tween 1% (TBST, Boston BioProducts, Cat: IBB-581X-4L) containing 3% BSA (Fisher, Cat: BP9703-100) for 10 minutes at room temperature (RT) while nutating. A 1:5,000 dilution of Streptavidin IR Dye 800CW (LI-COR, Cat: 926-32230) and/or a 1:5,000 dilution of rabbit anti-human PD-L1 mAb (Cell Signaling Technology, Cat: 13684S) were added to the membranes, followed by overnight incubation at 4°C while nutating.

The following day, membranes were washed 3x for 5 minutes with TBST and incubated with a 1:5,000 dilution of Streptavidin IR Dye 800CW (LI-COR, Cat: 926-32230) and/or a 1:7,100 dilution of IRDye 680RD goat anti-rabbit IgG secondary antibody (LI-COR, Cat: 926-68071) in 50ml of TBST for 1 hour at RT while nutating. The membranes were washed 3 x 5 minutes in TBST while nutating, then 2x with MilliQ-$H_2O$ and imaged on a LI-COR Odyssey CLx (LI-COR, Model 9140).

To evaluate total protein levels, the membranes were rinsed 3x with UltraPure distilled water (Invitrogen, Cat: 10977-015) and incubated in 20ml of GelCode™ Blue Stain Reagent (Thermo Fisher, Cat: 24590) for 5 minutes while nutating at RT. The membranes were washed 2x for 10 minutes each in 50ml of 50% methanol/1% acetic acid in UltraPure distilled water at RT while nutating. The membranes were rinsed 3x in UltraPure distilled water and imaged again on a LI-COR Odyssey CLx.

#### PD-L1-targeted Photolabeling on JY PD-L1 cells (Western Blot Analysis)

For two-antibody-based photolabeling, JY PD-L1 cells or JY WT cells (5 million cells per sample) were centrifuged for 5 minutes x 800xg, 4°C. Cells were resuspended in 4ml of cold 1x DPBS at a concentration of 5 million cells/ml and distributed into Protein LoBind microcentrifuge tubes in 1ml aliquots. Cells were pelleted by centrifugation (5 min x 800xg, 4°C) and resuspended in 1ml of 1x DPBS containing 5 µg Isotype control (BD Biosciences, Purified Mouse IgG1 κ, clone MOPC-21, Cat: 556648) or α-PD-L1 antibody (Invitrogen, Cat: 14-5983-82). Cells were incubated on a rotisserie for 30 min at 4°C and pelleted by centrifugation (5 min x 800xg, 4°C) to remove the supernatant. Pelleted cells were washed 2x with 1ml of cold 1x DPBS. Between washes, cells were pelleted by centrifugation (5 min x 800xg, 4°C). After the final wash, cells were resuspended in 1ml of cold 1x DPBS containing 5 µg of Goat α-Mouse IgG-RFT conjugate and incubated for 30 min at 4°C on a rotisserie. Following incubation, cells were centrifuged (5 min x

800xg, 4°C) and washed 2x with 1ml of cold 1x DPBS with centrifugation steps between washes (5 min x 800xg, 4°C). After the final wash, cells were resuspended in 1ml of cold 1x DPBS containing 250µM biotin phenol and irradiated at 100% light intensity for 0 min, 2 min or 10 min in a BPR200 bio-photoreactor. Cells were pelleted by centrifugation (5 min x 800xg, 4°C). Each cell pellet was resuspended in 150µl of a 1:1 mixture of RIPA Buffer and 4x Laemmli buffer supplemented with beta-mercaptoethanol per manufacturer's instructions (Fisher, Cat: BP176-100). Cells were sonicated 2x for 5 seconds at power level 6 and boiled for 5 minutes at 95°C. 15µl of each sample were loaded onto a Criterion TGX 12% Gel and subjected to western blotting procedures described above.

For blocking α-PD-L1 VHH-Fc-RFT-based photolabeling, JY PD-L1 cells (5 million cells per sample) were centrifuged for 5 minutes x 800xg, 4°C. Cells were resuspended in 4ml of cold 1x DPBS at a concentration of 5 million cells/ml and distributed into Protein LoBind microcentrifuge tubes in 1ml aliquots. Cells were pelleted by centrifugation and resuspended in 1ml of 1x DPBS containing 5µg of blocking α-PD-L1 VHH-Fc-RFT or Isotype VHH-Fc-RFT and incubated for 1 hour on a rotisserie at 4°C, then pelleted by centrifugation (5 min x 800xg, 4°C) to remove the supernatant. Pelleted cells were washed 2x with 1ml of 1x DPBS. Between washes, cells were pelleted by centrifugation (5 min x 800xg, 4°C), then resuspended in 500µl of 1x DPBS containing 250µM of biotin phenol and irradiated at 100% light intensity for 0 min, 2 min or 10 min in a BPR200 bio-photoreactor (Fisher Scientific: NC1558343) then centrifuged for 5 minutes at 800xg and 4°C. Each cell pellet was resuspended in 150µl of a 1:1 mixture of RIPA Buffer and 4x Laemmli buffer supplemented with beta-mercaptoethanol per manufacturer's instructions (Fisher, Cat: BP176-100). Cells were sonicated 2x for 5 seconds at power level 6 and boiled for 5 minutes at 95°C. 20µl of each sample were loaded onto a Criterion TGX 12% Gel and subjected to western blotting procedures described above.

#### Photolabeling of JY PD-L1-Jurkat JS86-PD-1 Co-Cultures

10 million JY PD-L1 or Jurkat JS86-PD-1 cells per VHH-Fc were centrifuged at 800xg for 5 minutes at 4°C and the cell pellet resuspended in 0.5ml cold complete media in Protein LoBind microcentrifuge tubes (1.5ml capacity, Eppendorf, Cat: 022431081). 5µg of the indicated VHH-Fcs were added per sample and incubated for 1 hour on a rotisserie at 4°C, followed by 2x washes with 1ml of 1x DPBS (with centrifugation at 800xg for 5 minutes at 4°C to pellet cells).

1 million VHH-targeted JY PD-L1 or Jurkat JS86-PD-1 cells were combined with 1 million untargeted JY PD-L1 or Jurkat JS86-PD-1 cells, spun down and the cell pellets resuspended in 45µl of 1x DPBS, followed by a 30-min co-culture at 37°C + 5% $CO_2$.

For RFT samples, 5µl of biotin phenol was added at a final concentration of 250µM and irradiated for 2 min at 100% light intensity in the biophotoreactor BPR200 (Efficiency Aggregators, Fisher, Cat: NC1558343). The samples were washed 1x with 1ml of 1x DPBS and centrifuged again.

For HRP samples, 5µl of reaction buffer (final concentration of 250 µM biotin phenol and 1 mM $H_2O_2$ in cold 1x DPBS) was added to each sample and incubated for 1 min at 4°C. The reactions were quenched with 50µl quenching buffer (5 mM Trolox, 10 mM Ascorbic Acid, and 10 mM $NaN_3$ in cold 1x DPBS), immediately pelleted by centrifugation (5 min at 800xg, 4°C), and washed again with 50µl of quenching buffer. The cells were then washed 1x with 1ml of 1x DPBS and centrifuged again. Samples were then processed for flow cytometry analysis or confocal microscopy analysis as indicated in the procedures below.

#### Flow Cytometry Analysis of Labeled Co-Cultures

Samples were photolabeled as described in Photolabeling of JY PD-L1-Jurkat JS86-PD-1 Co-Cultures. After light labeling, cells were washed 1x with 1ml of 1x DPBS and resuspended in 100µl of 1x DPBS containing BV421 Zombie Violet Fixable Viability Kit (BioLegend, Cat: 423113) at a 1:500 dilution, transferred to V-bottom plates (Falcon, Cat: 3894), and incubated for 15 minutes at room temperature, protected from light. The volume per well was brought up to 200µl with 1x DPBS and the cells were centrifuged for 5 minutes at 500xg, 4°C, washed 1x in 200µl DPBS and centrifuged again. Each cell pellet was resuspended in 100µl of Stain Buffer (BD Biosciences, Cat: 554656) with a 1:100 dilution of Fc Block (BD Biosciences, Cat: 564220) and incubated for 20 minutes on ice. The volume per well was brought up to 200µl with Stain Buffer and the cells were centrifuged for 5 minutes at 500xg, 4°C, washed 1x with 200µl Stain Buffer and centrifuged again. The following antibodies were used in 100µl of Stain Buffer per well: APC mouse anti-human CD19, clone HIB19 at a 1:10 dilution (BD Biosciences, Cat: 555415), Alexa Fluor 488 mouse anti-human CD3, clone SP34-2 at a 1:40 dilution (BD Biosciences, Cat: 557705), and Streptavidin PE at a 1:200 dilution (BD Biosciences, Cat: 554061). Compensation controls were prepared by adding 1 drop of UltraComp beads (BD Biosciences, Cat: 01-2222-42) to 100µl of Stain Buffer and 2µl of each individual antibody as each stained sample in FACS tubes (Fisherbrand, Cat: 14-956-3D), as well as PE mouse anti-human CD3, clone SK7 (BioLegend, Cat: 344806) and BV421 mouse anti-human CD45RO (BD Pharmingen, clone UCHL1, Cat: 555491) to set up PE and BV421 compensation controls, respectively. Both the experimental samples and the compensation control beads were stained for 30 minutes at 4°C.

The volume per sample was brought up to 200µl using Stain Buffer and then centrifuged for 5 minutes at 500xg, 4°C, washed 1x with 200µl of Stain Buffer, and centrifuged again. The cell pellets were resuspended in 200µl of Stain Buffer and transferred to 5ml FACS tubes. Samples were then acquired on a BD FACSCelesta with BD FACSDiva software. Data was analyzed using FlowJo, v10.

#### Confocal Microscopy Analysis of Labeled Monocultures or Co-Cultures

Samples were photolabeled as described in Photolabeling of JY PD-L1-Jurkat JS86-PD-1 Co-Cultures and prepared for confocal microscopy as in ref. 6. Briefly, microscope coverslips (Fisherbrand, Cat: 12-545-81) were acid-etched in 1N HCl (Fisher, Cat: SA56-1) for 30 minutes at 50°C, then washed in distilled water 3x and stored in 100% ethanol (Fisher, Cat: BP2818-500) at room temperature (RT). When ready to use, individual coverslips were placed into a 24-well plate (Thermo Fisher Scientific, Cat: 142485) and washed 2x with 1ml of 1x DPBS (Gibco, Cat: 14190-144). Coverslips were incubated in 500µl of poly-L-lysine solution (Sigma, Cat: P4707-50ML) for 30 minutes at 37°C, then washed 2x with 1ml of 1x DPBS. 500µl of labeled cells were loaded per well and centrifuged for 3 minutes at 400xg and 4°C with deceleration set at 3 using a Sorvall Legend XTR table-top centrifuge (Thermo Scientific).

Supernatants were removed and samples were fixed in 1ml of 1x cold DPBS containing a final concentration of 3% paraformaldehyde (PFA, Electron Microscopy Sciences, Cat: 15710) and 0.1% glutaraldehyde (Sigma-Aldrich, Cat: G5882-10) for 10 minutes at 4°C. Coverslips were washed 3x in 1ml of Stain Buffer (BD Biosciences, Cat: 554656) and incubated overnight at 4°C in 1ml of Stain Buffer.

The following day, labeled monoculture samples were stained with a 1:200 dilution of Alexa Fluor 488 Streptavidin (BioLegend, Cat: 405235) in 500µl Stain Buffer. Co-culture samples were stained with a 1:200 dilution of Alexa Fluor 488 Streptavidin and a 1:40 dilution of BV650 Mouse Anti-Human CD3 antibody (BD Biosciences, Cat: 563999) in 500µl of Stain Buffer. Plates were sealed with parafilm and incubated overnight at 4°C, protected from light.

Samples were washed 1x in 1ml of Stain Buffer and Hoechst DNA dye (Cayman Chemical Company, Cat: 600332) was added at a 1:10,000 dilution in 500µl of Stain Buffer and incubated for 10 minutes at RT, protected from light. Samples were washed 2x in 1ml of Stain Buffer and fixed with 500µl of 1x DPBS containing 3% PFA and 0.1% glutaraldehyde for 5 minutes at RT. Coverslips were washed 2x with 1ml of Stain Buffer, then mounted using a drop of ProLong Gold Anti-fade mountant (Invitrogen, Cat: P36934) on microscope slides (J. Melvin Freed, Cat: 301MF). Mounted samples were dried overnight at RT, protected from light. Images were collected on a Zeiss LSM800 confocal microscope using Airyscan settings at 63X magnification and ZEN 2 software (Zeiss).

#### RFT-based Photoproximity Labeling of Proteins for LC-MS/MS Analysis

For two-antibody-based RFT photoproximity labeling in monoculture, Jurkat NF-κβ GFP or JY-PD-L1 cells (20 or 50 million cells per sample) were transferred to 50ml conical tubes and pelleted by centrifugation (5 min at 800xg, 4$^{o}$C). The cells were then resuspended in cold 1x DPBS and combined to a single conical tube and pelleted by centrifugation (5 min at 800xg, 4$^{o}$C). The supernatant was removed, and the cells were resuspended at a concentration of 20 or 50 million cells/ml in cold 1x DPBS. The cell suspension was transferred to Protein LoBind tubes in 1ml experimental aliquots and pelleted by centrifugation (4 min at 1,000xg, 4$^{o}$C). For each experiment, cells from each aliquot were resuspended in 1ml of cold 1x DPBS containing isotype control (BD Biosciences Purified Mouse IgG1 κ, clone MOPC-21: 556648) or α-PD-L1 antibody (Thermo Fisher Scientific: 14598382) for JY PD-L1 cells. 0.5 µg of primary antibody was added for every 10 million cells (10 or 25µg per sample). The cells were incubated on a rotisserie for 30 min at 4$^{o}$C. After incubation, the cells were pelleted by centrifugation (4 min at 1,000xg, 4$^{o}$C) to remove the supernatant. The cells were then washed 2x with 1ml cold 1x DPBS. After the final wash, the cells were pelleted by centrifugation (4 min at 1,000xg, 4$^{o}$C) and resuspended in 1ml of cold 1x DPBS containing 10 or 25µg of Goat α-Mouse IgG-RFT conjugate (prepared as described above) and incubated on a rotisserie for 30 min at 4$^{o}$C. After incubation, the cells were pelleted by centrifugation (4 min at 1,000xg, 4$^{o}$C) to remove the supernatant. The cells were then washed 2x with 1ml cold 1x DPBS. After the final wash, the cells were pelleted by centrifugation (4 min at 1,000xg, 4$^{o}$C) and resuspended in 1ml of cold 1x DPBS containing 250µM biotin phenol. The samples were placed in the bio-photoreactor for 2 min or 10 min and irradiated at full intensity.

For VHH-Fc-RFT-based photoproximity labeling in monoculture, JY PD-L1 cells (20 million cells per sample) were transferred to 50ml conical tubes and pelleted by centrifugation (5 min at 800xg, 4$^{o}$C). The cells were then resuspended in cold 1x DPBS and combined to a single conical tube and pelleted by centrifugation (5 min at 800xg, 4$^{o}$C). The supernatant was removed, and the cells were resuspended at a concentration of 20 million cells/ml in cold 1x DPBS. The cell suspension was transferred to Protein LoBind tubes in 1ml experimental aliquots and pelleted by centrifugation (4 min at 1,000xg, 4$^{o}$C). For each experiment, cells from each aliquot were resuspended in 1ml of cold 1x DPBS containing 10µg of Isotype VHH-Fc-RFT, blocking α-PD-L1 VHH-Fc-RFT, or non-blocking α-PD-L1 VHH-Fc-RFT (prepared as described above). The cells were incubated on a rotisserie for 1 hr at 4$^{o}$C. After incubation, the cells were pelleted by centrifugation (4 min at 1,000xg, 4$^{o}$C) to remove the supernatant. The cells were then washed 2x with 1ml cold 1x DPBS. After the final wash, the cells were pelleted by centrifugation (4 min at 1,000xg, 4$^{o}$C) and resuspended in 1ml of cold 1x DPBS containing 250µM biotin phenol. The samples were placed in the bio-photoreactor for 2 min and irradiated at full intensity.

For VHH-Fc-RFT-based photoproximity labeling within JY PD-L1-Jurkat JS86-PD-1 immune synapses, JY PD-L1 or Jurkat JS86-PD-1 cells (20 million cells per sample) were centrifuged at 800xg for 5 minutes at 4°C and resuspended in cold complete medium at a concentration of 20 million cells/ml. The cell suspension was transferred to Protein LoBind tubes in 1ml experimental aliquots. For each experiment, 10µg of

Isotype VHH-Fc-RFT for either JY PD-L1 or Jurkat JS86-PD-1 cells, non-blocking α-PD-L1 VHH-Fc-RFT for JY PD-L1 cells only, or non-blocking α-PD-1 VHH-Fc-RFT for Jurkat JS86-PD-1 cells only (prepared as described above) was added to the aliquots. The cells were incubated on a rotisserie for 1 hr at 4$^o$C. After incubation, the cells were pelleted by centrifugation (4 min at 1,000xg, 4$^o$C) to remove the supernatant. The cells were then washed 2x with 1ml cold 1x DPBS. After the final wash, 20 million VHH-Fc-RFT targeted JY PD-L1 or Jurkat JS86-PD-1 cells were combined with 20 million non-targeted Jurkat JS86-PD-1 cells or JY PD-L1 cells, respectively and centrifuged for 4 min at 800xg, 4$^o$C. Pelleted cells were gently resuspended in 900µl of cold 1x DPBS and incubated for 30 min at 37$^o$C. Following incubation, 100µl of 2.5mM biotin phenol (final concentration 250µM) was added and samples were irradiated for 2 min at full intensity in the bio-photoreactor.

After irradiation, cells were then pelleted by centrifugation (4 min at 1,000xg, 4$^o$C) to remove the supernatant. The cells were then washed 2x with 1ml cold 1x DPBS. After the final wash, the cells were pelleted by centrifugation (4 min at 1,000xg, 4$^o$C) and resuspended in 1ml of membrane permeabilization buffer (MEM-PER Plus Membrane Fractionation Kit, Thermo Fisher Scientific: 89842) containing 1x protease inhibitors (Sigma-Aldrich: 11873580001) and incubated for 20 min at 4$^o$C. The samples were then spun at 16,000xg for 15 min at 4$^o$C. The supernatant containing cytosolic proteins was removed, and the pellet was resuspended in 300µl lysis buffer (RIPA buffer, Thermo Fisher Scientific: 89900) containing 1% SDS and 1x protease inhibitors. The samples were sonicated to break up the membrane pellet (1x 5s at power level 6) and then heated for 5 min at 95$^o$C. The samples were then diluted to 1.3ml with RIPA and sonicated (2x5s at power level 6). The protein concentration was measured by BCA and the samples were stored at -80$^o$C till streptavidin bead enrichment. For bead enrichment, 1mg of cell lysate was added to a Protein LoBind tube containing 250µl of streptavidin magnetic beads (Thermo Fisher Scientific: 88817) that were pre-washed 2x with 1ml RIPA buffer. The samples were incubated for 3 hours at room temperature, and the beads were pelleted on a magnetic rack. The supernatant was removed, and the beads were washed with 3x 1ml 1%SDS in 1x DPBS, 3x 1ml 1M NaCl in 1x DPBS, 3x 1ml 10% EtOH in 1x DPBS, and once in 1ml RIPA. The beads were incubated in each of the washes for 5 min before pelleting. After the final wash, the beads were resuspended in 30µl of 4x Laemmli sample buffer containing 20mM DTT and 25mM Biotin and heated to 95$^o$C for 10 min. The samples were placed on the magnetic rack and the supernatant was collected and transferred to a new Protein LoBind tube and stored at -80$^o$C until quantitative proteomic sample preparation and analysis (performed at IQ Proteomics, Cambridge, MA).

For untargeted labeling of Jurkat NF-κβ GFP cells, 300 million Jurkat NF-κβ GFP cells growing at 1 million cells/ml were removed from cell culture and transferred to 50ml conical vials and pelleted by centrifugation (5 min at 800xg, 4$^o$C). The cells were then resuspended in cold 1x DPBS and recombined to a single conical vial and pelleted by centrifugation (5 min at 800xg, 4$^o$C). The supernatant was removed, and the cells were resuspended in 6ml of PBS at 50 million cells/ml. The cell suspension was transferred to Protein LoBind tubes in 1ml aliquots and pelleted by centrifugation (4 min at 1,000xg, 4$^o$C). The cells were then resuspended in 1ml of 50µM RFT alkyne plus 250µM biotin phenol, or 250µM biotin phenol only in DPBS. The samples were then irradiated for 10 min in the bio-photoreactor at 100% intensity. After irradiation, the cells were pelleted by centrifugation (4 min at 1,000xg, 4$^o$C) to remove the supernatant. The cells were then washed 2x with 1ml cold 1x DPBS. After the final wash, the cells were pelleted by centrifugation (4 min at 1,000xg, 4$^o$C). The supernatant was removed, the cell pellets were lysed, and the biotinylated proteins were enriched from the lysate according to the streptavidin enrichment and elution steps described above.

#### VHH-Fc-HRP Photoproximity Labeling within JY PD-L1-Jurkat JS86-PD-1 Immune Synapses for LC-MS/MS Analysis

JY PD-L1 cells (20 million cells per sample) were centrifuged at 800xg for 5 minutes at 4°C and resuspended in cold complete medium at a concentration of 20 million cells/ml. The cell suspension was transferred to Protein LoBind tubes in 1ml experimental aliquots. For each experiment, 10µg of Isotype VHH-Fc-HRP or non-blocking α-PD-L1 VHH-Fc-HRP (prepared as described above) was added to the aliquots. The cells were incubated on a rotisserie for 1 hr at 4ºC. After incubation, the cells were pelleted by centrifugation (4 min at 1,000xg, 4ºC) to remove the supernatant. The cells were then washed 2x with 1ml cold 1x DPBS. After the final wash, 20 million VHH-Fc-HRP targeted JY PD-L1 cells were combined with 20 million non-targeted Jurkat JS86-PD-1 cells and centrifuged for 4 min at 800xg, 4ºC. Pelleted cells were gently resuspended in 900µl of cold 1x DPBS and incubated for 30 min at 37ºC. Following incubation, the sample volume was brought to 1ml with cold 1x DPBS and added directly to a 15 ml conical tube containing 4ml of reaction buffer (250µM biotin phenol and 1mM $H_2O_2$ in cold 1x DPBS) and incubated for 1 min at 4ºC. After 1 min, 5ml quenching buffer (1x DPBS containing 5mM Trolox, 10mM Sodium Ascorbic Acid, 10mM $NaN_3$) was added to the cells. The cells were then pelleted by centrifugation (4 min at 1,000xg, 4ºC), washed once with 10ml of quenching buffer (1x DPBS containing 5mM Trolox, 10mM Sodium Ascorbic Acid, 10mM $NaN_3$), resuspended in 1ml of cold 1x DPBS and transferred to Protein LoBind tubes. The cells were then washed 2x in cold 1x DBPS, and pelleted by centrifugation (4 min at 1,000xg, 4ºC). Cell pellets were resuspended in 1ml of membrane permeabilization buffer (MEM-PER Plus Membrane Fractionation Kit, Thermo Fisher Scientific: 89842) containing 1x protease inhibitors (Sigma-Aldrich: 11873580001) and incubated for 20 min at 4ºC. The samples were then spun at 16,000xg for 15 min at 4ºC. The supernatant containing cytosolic proteins was removed, and the pellet was resuspended in 300µl lysis buffer (RIPA buffer, Thermo Fisher Scientific: 89900) containing 1% SDS and 1x protease inhibitors. The samples were sonicated to break up the membrane pellet (1x 5s at power level 6) and then heated for 5 min at 95ºC. The samples were then diluted to 1.3ml with RIPA and sonicated (2x5s at power level 6). The protein concentration was measured by BCA and the samples were stored at -80ºC till streptavidin bead enrichment. For bead enrichment, 1mg of cell lysate was added to a Protein LoBind tube containing 250µl of streptavidin magnetic beads (Thermo Fisher Scientific: 88817) that were pre-washed 2x with 1ml RIPA buffer. The samples were incubated for 3 hours at room temperature, and the beads were pelleted on a magnetic rack. The supernatant was removed, and the beads were washed with 3x 1ml 1%SDS in 1x DPBS, 3x 1ml 1M NaCl in 1x DPBS, 3x 1ml 10% EtOH in 1x DPBS, and once in 1ml RIPA. The beads were incubated in each of the washes for 5 min before pelleting. After the final wash, the beads were resuspended in 30µl of 4x Laemmli sample buffer containing 20mM DTT and 25mM Biotin and heated to 95ºC for 10 min. The samples were placed on the magnetic rack and the supernatant was collected and transferred to a new Protein LoBind tube and stored at -80ºC until quantitative proteomic sample preparation and analysis (performed at IQ Proteomics, Cambridge, MA).

#### VHH-Fc-Ir Photoproximity Labeling within JY PD-L1-Jurkat JS86-PD-1 Immune Synapses for LC-MS/MS Analysis

JY PD-L1 cells (20 million cells per sample) were centrifuged at 800xg for 5 minutes at 4°C and resuspended in cold complete medium at a concentration of 20 million cells/ml. The cell suspension was transferred to Protein LoBind tubes in 1ml experimental aliquots. For each experiment, 10µg of Isotype VHH-Fc-Ir or non-blocking α-PD-L1 VHH-Fc-Ir (prepared as described above) was added to the aliquots. The cells were incubated on a rotisserie for 1 hr at 4ºC. After incubation, the cells were pelleted by centrifugation (4 min at 1,000xg, 4ºC) to remove the supernatant. The cells were then washed 2x with 1ml cold 1x DPBS. After the final wash, 20 million VHH-Fc-Ir targeted JY PD-L1 cells were combined with 20 million non-targeted Jurkat JS86-PD-1 cells and centrifuged for 4 min at 800xg, 4ºC. Pelleted cells were gently resuspended in

900µl of cold 1x DPBS and incubated for 30 min at 37$^{o}$C. Following incubation, 100µl of 2.5mM Dz (250µM final concentration) was added and samples were irradiated for 2 min at full intensity in the bio-photoreactor. After irradiation, cells were then pelleted by centrifugation (4 min at 1,000xg, 4$^{o}$C) to remove the supernatant. The cells were then washed 2x with 1ml cold 1x DPBS. After the final wash, the cells were pelleted by centrifugation (4 min at 1,000xg, 4$^{o}$C) and resuspended in 1ml of membrane permeabilization buffer (MEM-PER Plus Membrane Fractionation Kit, Thermo Fisher Scientific: 89842) containing 1x protease inhibitors (Sigma-Aldrich: 11873580001) and incubated for 20 min at 4$^{o}$C. The samples were then spun at 16,000xg for 15 min at 4$^{o}$C. The supernatant containing cytosolic proteins was removed, and the pellet was resuspended in 300µl lysis buffer (RIPA buffer, Thermo Fisher Scientific: 89900) containing 1% SDS and 1x protease inhibitors. The samples were sonicated to break up the membrane pellet (1x 5s at power level 6) and then heated for 5 min at 95$^{o}$C. The samples were then diluted to 1.3ml with RIPA and sonicated (2x5s at power level 6). The protein concentration was measured by BCA and the samples were stored at -80$^{o}$C till streptavidin bead enrichment. For bead enrichment, 1mg of cell lysate was added to a Protein LoBind tube containing 250µl of streptavidin magnetic beads (Thermo Fisher Scientific: 88817) that were pre-washed 2x with 1ml RIPA buffer. The samples were incubated for 3 hours at room temperature, and the beads were pelleted on a magnetic rack. The supernatant was removed, and the beads were washed with 3x 1ml 1%SDS in 1x DPBS, 3x 1ml 1M NaCl in 1x DPBS, 3x 1ml 10% EtOH in 1x DPBS, and once in 1ml RIPA. The beads were incubated in each of the washes for 5 min before pelleting. After the final wash, the beads were resuspended in 30µl of 4x Laemmli sample buffer containing 20mM DTT and 25mM Biotin and heated to 95$^{o}$C for 10 min. The samples were placed on the magnetic rack and the supernatant was collected and transferred to a new Protein LoBind tube and stored at -80$^{o}$C until quantitative proteomic sample preparation and analysis (performed at IQ Proteomics, Cambridge, MA).

#### Protein Extraction and Digestion for LC-MS/MS Analysis

Prior to LC-MS/MS analysis, proteins were reduced with 20 mM dithiothreitol (DTT) for 1 hour at room temperature. Cysteine residues were alkylated with iodoacetamide (60 mM) for 1 hour in the dark and quenched with DTT (40 mM). Proteins were extracted by methanol-chloroform precipitation and digested with 1 µg of trypsin (Promega) in 100 mM EPPS (pH 8.0) for 4 hours at 37°C. Each of the tryptic peptide samples were labeled with 400 µg of Tandem Mass Tag (TMT; Pierce) isobaric reagents for 2 hours at room temperature. A label efficiency check was performed by pooling 2µl from each sample within a single plex to ensure at least 98% labeling of all N-termini and lysine residues. All samples were quenched with hydroxylamine (0.5%), acidified with TFA (2%), pooled, and dried by speedvac evaporation. Pooled TMT labeled peptides were fractionated using the high pH reverse-phase peptide fractionation kit (Pierce) into 3 fractions (20%, 25%, and 50% acetonitrile in 0.1% triethylamine) and desalted with Empore-C18 (3M) in-house packed StageTips prior to analysis by mass spectrometry.

#### LC-MS/MS-based Proteomic Analysis of Labeled Cell Experiments

All mass spectra were acquired on an Orbitrap Fusion Lumos coupled to an EASY nanoLC-1000 (or nanoLC-1200) (Thermo Fisher) liquid chromatography system. Approximately 2 µg of peptides were loaded on a 75 µm capillary column packed in-house with Sepax GP-C18 resin (1.8 µm, 150 Å, Sepax) to a final length of 35 cm. Peptides were separated using a 110-minute linear gradient from 8% to 28% acetonitrile in 0.1% formic acid. The mass spectrometer was operated in a data dependent mode. The scan sequence began with FTMS1 spectra (resolution = 120,000; mass range of 350-1400 *m/z*; max injection time of 50 ms; AGC target of 1e6; dynamic exclusion for 60 seconds with a +/- 10 ppm window). The ten most intense precursor ions were selected for MS2 analysis via collisional-induced dissociation (CID) in the ion trap (normalized collision energy (NCE) = 35; max injection time = 100ms; isolation window of 0.7 Da; AGC target of 1.5e4). A real-time search (RTS) approach was utilized during data acquisition to only trigger quantitative spectra on high-confidence peptide identifications[7]. Online spectral identification was accomplished via a custom software client that monitors spectral acquisition though a vendor supplied instrument application programming interface and assigns peptide sequences through a probabilistic model, in real-time (Orbitrap Lumos). The RTS client utilized a peptide database composed of in-silico predicated human tryptic peptides. Confidently identified peptides were quantified via a synchronous-precursor-selection (SPS) MS3 method that selected eight MS2 product ions for high energy collisional-induced dissociation (HCD) with analysis in the Orbitrap (NCE = 55; resolution = 50,000; max injection time = 200 ms; AGC target of 1.4e5; isolation window at 1.2 Da for +2 *m/z*, 1.0 Da for +3 *m/z* or 0.8 Da for +4 to +6 *m/z*). All mass spectra were converted to mzXML using a modified version of ReAdW.exe. MS/MS spectra were searched against a concatenated 2021 human Uniprot protein database containing common contaminants (forward + reverse sequences) using the SEQUEST algorithm[8]. Database search criteria are as follows: fully tryptic with two missed cleavages; a precursor mass tolerance of 50 ppm and a fragment ion tolerance of 1 Da; oxidation of methionine (15.9949 Da) was set as differential modifications. Static modifications were carboxyamidomethylation of cysteines (57.0214) and TMT on lysines and N-termini of peptides (229.1629). Peptide-spectrum matches were filtered using linear discriminant analysis[9] and adjusted to a 1% peptide false discovery rate (FDR)[10].

#### Bioinformatic Analysis of LC-MS/MS Data

All bioinformatic analysis of LC-MS/MS data was performed in the R statistical computing environment[11]. Peptide level abudance data is used to identify the number of peptides corresponding to a protein in the experiment. Any protein with a single peptide quantification is removed to reduce the possibility that outliers will affect downstream proximal calls. Peptide level abundance data is then normalized to the summed total abundance for each sample separately. These totals are then averaged, and each normalized protein abundance value is multiplied by this average to rescale abundance data. Peptide level data is then merged to protein level data by taking the median of all peptides corresponding to a protein.

#### Linear Modeling and Fold Change Generation

For experiments in which only two conditions are tested, protein abundances are $\log_2$ transformed and subjected to linear modeling analysis with Limma[12]. Limma utilizes an empirical Bayes approach that allows for a realistic distribution of biological variance with small sample sizes per group. This program further utilizes the full dataset to shrink the observed sample variances towards a pooled estimate. This borrowing of variance information across proteins allows for a more accurate estimate of true variance, and improved power to detect real differences between groups. For each protein, abundance data is fit to a linear model using the lmFit function with the experimental group set as the covariate. The $\log_2$FC values are estimated and p-values calculated for significance. P-values were also corrected for multiple comparisons using the false discovery rate (FDR) method by Benjamini and Hochberg[13].

#### Volcano Plot Generation

Volcano plots were generated in R with the ggplot library[14]. $\text{Log}_2$FC and p-value estimates from Limma were subset to those reaching a $\log_2$FC described in the figure legends for each experiment. Proteins were colored based on whether they fell above or below the $\log_2$-fold cutoff threshold and were statistically significant (p-value of $< 0.05$).

#### STRING Plot Analysis

Proteins reaching significance and the $\log_2$FC cutoff defined in the experiment were analyzed with StringDB for associations between proteins to generate a protein interaction map. Network data was downloaded from the StringDB analysis and imported into Cytoscape 3.8.2[15] for visualization. Edges in this network indicate interaction from all sources within the StringDB database; textmining, experiments, databases, co-expression, neighborhood, gene fusion, and co-occurrence. The minimum required overall interaction confidence score for an edge was set to 0.4, with only query proteins shown. Thick edges are supported by evidence from the "experiments" category from StringDB with a confidence level above 0.1, and thin edges represent a connection with evidence from other categories. Gene set enrichment analysis was performed within StringDB and used to place nodes into broad GO (biological process) terms indicated by coloration defined in each figure. A pie graph node is utilized for proteins belonging to multiple groups.

#### Tumor (TCGA) vs Normal (GTEx) Separation Score Analysis

A combined cohort of TCGA and GTEx RNA expression data was obtained from the UCSC Xena portal (RSEM tpm (n=19,131) UCSC Toil RNA-seq Recompute (UCSC Xena (xenabrowser.net)[16]. Tumor data was subset to primary disease samples only. Tumor and normal samples were paired by the provided primary site annotation. The separation score was adapted from prior work[17] which explores tumor and normal samples as clusters in unified gene expression space. We utilized metrics described in the publication: the log normalized Davies-Bouldin score, a metric that measures the ratio of within tumor to normal sample cluster spread to cluster distance; and the Manhattan distance between the centroids of tumor and normal co-expression for two genes. We then subtracted the log-transformed normalized DB score from log-transformed Manhattan distance score to arrive at a separation score. The higher separation scores represent better gene expression segregation of tumor to normal samples for pairs of genes. Pairs with high segregation provide opportunities for development of targeted therapies with boolean gate approaches, including Antigen 1 AND Antigen 2, Antigen 1 AND NOT Antigen 2, or Antigen 2 AND NOT Antigen 1 logic gates.

Data analysis and visualization were performed with R statistical programming language (4.1.0) and tidyverse (1.3.1). Heatmaps were generated using pheatmap package (1.0.12) and the venn diagram was generated with VennDiagram package (1.7.0).

#### Tumor (TCGA) Correlation Analysis

A combined cohort of TCGA and GTEx RNA expression data was obtained from the UCSC Xena portal (RSEM tpm (n=19,131) UCSC Toil RNA-seq Recompute and subset to primary pancreatic tumor samples only. Pearson correlation estimate is computed for genes and visualized using corrplot R package (0.84). Correlations between gene pairs for which the estimates fail significance cutoff at $p > 0.05$ are colored in white.

#### Single Cell RNAseq

The visualized single cell RNAseq data is a collection of previously published datasets and was obtained from BioTuring Database (access date 12/15/2021). At time of access the resource comprises 255 studies with a total of 14,031,384 cells used to generate the visualization. The dot plot of genes across cell types of interest was generated using Talk2Data Bioturing feature with Vinci software (BioTuring Inc., San Diego, CA, USA). The dot size represents percentage of cells with expression, and color corresponds to the average rank of expression. All data is represented on a relative scale.

#### Gene Ontology Enrichment

Gene set enrichment analysis of GO Biological Process (GO_Biological_Process_2021 library) was conducted in R using enrichR (2.1)[18]. The reported log-scaled p-value was computed with Fisher's exact test with Benjamini-Hochberg multiple hypothesis correction.

### Supplementary References

1. UniProt, C. UniProt: the universal protein knowledgebase in 2021. *Nucleic Acids Res* **49**, D480-D489 (2021).
2. Bausch-Fluck, D. et al. The in silico human surfaceome. *Proc Natl Acad Sci U S A* **115**, E10988-E10997 (2018).
3. Geri, J.B. et al. Microenvironment mapping via Dexter energy transfer on immune cells. *Science* **367**, 1091-+ (2020).
4. Szklarczyk, D. et al. STRING v11: protein-protein association networks with increased coverage, supporting functional discovery in genome-wide experimental datasets. *Nucleic Acids Res* **47**, D607-d613 (2019).
5. Shao, X. et al. CellTalkDB: a manually curated database of ligand-receptor interactions in humans and mice. *Briefings in Bioinformatics* **22**(2021).
6. Oslund, R.C. et al. Detection of cell-cell interactions via photocatalytic cell tagging. *Nat Chem Biol* **18**, 850-858 (2022).
7. Erickson, B.K. et al. Active Instrument Engagement Combined with a Real-Time Database Search for Improved Performance of Sample Multiplexing Workflows. *Journal of Proteome Research* **18**, 1299-1306 (2019).
8. Eng, J.K., Mccormack, A.L. & Yates, J.R. An Approach to Correlate Tandem Mass-Spectral Data of Peptides with Amino-Acid-Sequences in a Protein Database. *Journal of the American Society for Mass Spectrometry* **5**, 976-989 (1994).
9. Huttlin, E.L. et al. A Tissue-Specific Atlas of Mouse Protein Phosphorylation and Expression. *Cell* **143**, 1174-1189 (2010).
10. Elias, J.E. & Gygi, S.P. Target-decoy search strategy for increased confidence in large-scale protein identifications by mass spectrometry. *Nature Methods* **4**, 207-214 (2007).
11. R Development Core Team, R. R: A language and environment for statistical computing. (R foundation for statistical computing Vienna, Austria, 2011).
12. Ritchie, M.E. et al. limma powers differential expression analyses for RNA-sequencing and microarray studies. *Nucleic Acids Research* **43**, e47-e47 (2015).
13. Benjamini, Y. & Hochberg, Y. Controlling the False Discovery Rate: A Practical and Powerful Approach to Multiple Testing. *Journal of the Royal Statistical Society. Series B (Methodological)* **57**, 289-300 (1995).
14. Wickham, H. *ggplot2: Elegant Graphics for Data Analysis*, (Springer-Verlag, New York, 2016).
15. Shannon, P. et al. Cytoscape: a software environment for integrated models of biomolecular interaction networks. *Genome Res* **13**, 2498-504 (2003).
16. Goldman, M.J. et al. Visualizing and interpreting cancer genomics data via the Xena platform. *Nature Biotechnology* **38**, 675-678 (2020).
17. Dannenfelser, R. et al. Discriminatory Power of Combinatorial Antigen Recognition in Cancer T Cell Therapies. *Cell Systems* **11**, 215-+ (2020).
18. Kuleshov, M.V. et al. Enrichr: a comprehensive gene set enrichment analysis web server 2016 update. *Nucleic Acids Res* **44**, W90-7 (2016).
